# Multimodal CRISPR screens uncover DDX39B as a global repressor of A-to-I RNA editing

**DOI:** 10.1101/2024.11.22.624788

**Authors:** Tianzi Wei, Jiaxuan Li, Xiang Lei, Risheng Lin, Qingyan Wu, Zhenfeng Zhang, Shimin Shuai, Ruilin Tian

**Affiliations:** Department of Medical Neuroscience, School of Medicine, Southern University of Science and Technology, Shenzhen, Guangdong 518055, China; Key University Laboratory of Metabolism and Health of Guangdong, Southern University of Science and Technology, Shenzhen, Guangdong 518055, China; Department of Human Cell Biology and Genetics, School of Medicine, Southern University of Science and Technology, Shenzhen, Guangdong 518055, China; School of Public Health and Emergency Management, Southern University of Science and Technology, Shenzhen, Guangdong 518055, China

**Author notes:** These authors contributed equally (Tianzi Wei, Jiaxuan Li, Xiang Lei). Correspondence (S.S.), (R.T.).

## Abstract

Adenosine-to-Inosine (A-to-I) RNA editing is a critical post-transcriptional modification that diversifies the transcriptome and influences various cellular processes. Despite its significance, the regulatory mechanisms controlling A-to-I editing remain largely unknown. In this study, we present two complementary CRISPR-based genetic screening platforms: CREDITS (<u>C</u>RISPR-based <u>R</u>NA <u>EDIT</u>ing regulator <u>S</u>creening), which enables genome-scale identification of editing regulators using an RNA recorder-based reporter system, and scCREDIT-seq (<u>s</u>ingle-<u>c</u>ell <u>C</u>RISPR-based <u>R</u>NA <u>EDIT</u>ing <u>seq</u>uencing), which provides multiplexed single-cell characterization of transcriptome and editome changes for pooled perturbations on dozens of selected genes. Through screening 1,350 RNA-binding proteins, we identified a series of known and novel A-to-I editing regulators. Detailed mechanistic investigation revealed DDX39B as a global repressor of A-to-I editing, which functions by preventing double-stranded RNA accumulation through its helicase and ATPase activities. We demonstrate that targeting DDX39B significantly enhances the efficiency of RNA editing-based tools like CellREADR and LEAPER, and represents a promising strategy for anti-HDV therapy by modulating viral genome editing. These technological advances not only expand our understanding of RNA editing regulation but also provide powerful tools for exploring tissue-specific and context-dependent RNA modification mechanisms, with broad implications for therapeutic development.

## Introduction

Adenosine-to-Inosine (A-to-I) editing is a prevalent post-transcriptional RNA modification, in which ADAR (Adenosine Deaminase Acting on RNA) enzymes convert adenosine to inosine through hydrolytic deamination. Inosine is recognized as guanosine by cellular machinery, effectively recoding genetic information and enhancing transcriptome diversity^1–4^.

The ADAR family comprises three main proteins in mammals: ADAR1 (*ADAR*), which exists in two isoforms (p150 and p110), ubiquitously expressed in all tissues; ADAR2 (*ADARB1*), primarily expressed in the brain; and ADAR3 (*ADARB2*), which is catalytically inactive but may have regulatory roles. These enzymes target double-stranded RNA (dsRNA) regions formed by inverted repeat sequences, complementary sequences in adjacent introns, or complex RNA secondary structures^5–7^.

Recent advances in high-throughput sequencing technologies have led to the identification of millions of A-to-I editing sites (the editome) across the human transcriptome, revealing its widespread occurrence in both coding and non-coding regions^8–12^. In coding sequences, editing can lead to amino acid recoding, potentially altering protein function. In non-coding regions, editing can affect RNA splicing, stability, and localization.

The precise spatiotemporal regulation of A-to-I editing is crucial for maintaining cellular function and homeostasis, as exemplified in neural function and immune response regulation. In the nervous system, editing modifies neurotransmitter receptors, regulates synaptic transmission, and influences neural development^13–16^. Within the immune system, A-to-I editing plays crucial roles in viral RNA modification, innate immunity regulation, and the discrimination between self and non-self RNA^17–20^. Dysregulation of RNA editing has been associated with numerous diseases, such as cancer, neurological disorders, and autoimmune diseases^21–25^.

Recent efforts have identified additional A-to-I regulators beyond the well-characterized ADAR proteins^8,26–31^. Yet, the complex regulatory network controlling editing efficiency and specificity remains poorly understood. Discovering new A-to-I regulators holds promise for advancing our understanding of RNA editing mechanisms, improving RNA editing-based technologies for programmable RNA modifications and targeted cell manipulation, and developing therapeutics for related diseases.

Here, we established high-throughput CRISPR-based genetic screening platforms that enable systematic identification and multiplexed transcriptome and editome characterization of A-to-I RNA editing regulators. Through these technologies, we uncovered multiple previously unknown regulators of A-to-I editing and elucidated the role of *DDX39B* as a global repressor of A-to-I editing. We further demonstrated targeting DDX39B as a potential strategy to enhance RNA editing-based tools and develop antiviral therapies.

## Results

### Development of CREDITS for pooled CRISPR screening on A-to-I editing

To enable pooled CRISPR screening for A-to-I editing regulators, we developed a method called CREDITS (<u>C</u>RISPR-based <u>R</u>NA <u>EDIT</u>ing regulator <u>S</u>creening). The general principle of this method is to associate sgRNA-induced genetic perturbations with A-to-I editing outcomes of an RNA editing recorder. We reasoned that an RNA fragment that contains known editing sites in a dsRNA structure could serve as a molecular recorder for A-to-I editing events. One of the well-established A-to-I editing sites in human cells is the Q/R conversion site in *GRIA2* gene^14^. We validated the endogenous editing of this site in a neuroblastoma cell line SH-SY5Y and in iPSC-derived neurons (Fig. 1a), consistent with previous reports that the Q/R conversion event of *GRIA2* is catalyzed by ADAR2 which is expressed mostly in neuronal cells^15,32^. Therefore, a fragment of *GRIA2* containing the Q/R conversion site and the flanking sequences predicted to form a dsRNA structure was used as a candidate RNA editing recorder for further characterization (Fig. 1b). The editing outcomes of the recorder can be retrieved by Sanger sequencing or NGS, as A-to-I editing events lead to A-to-G conversions in RT-PCR.

**Fig. 1:**
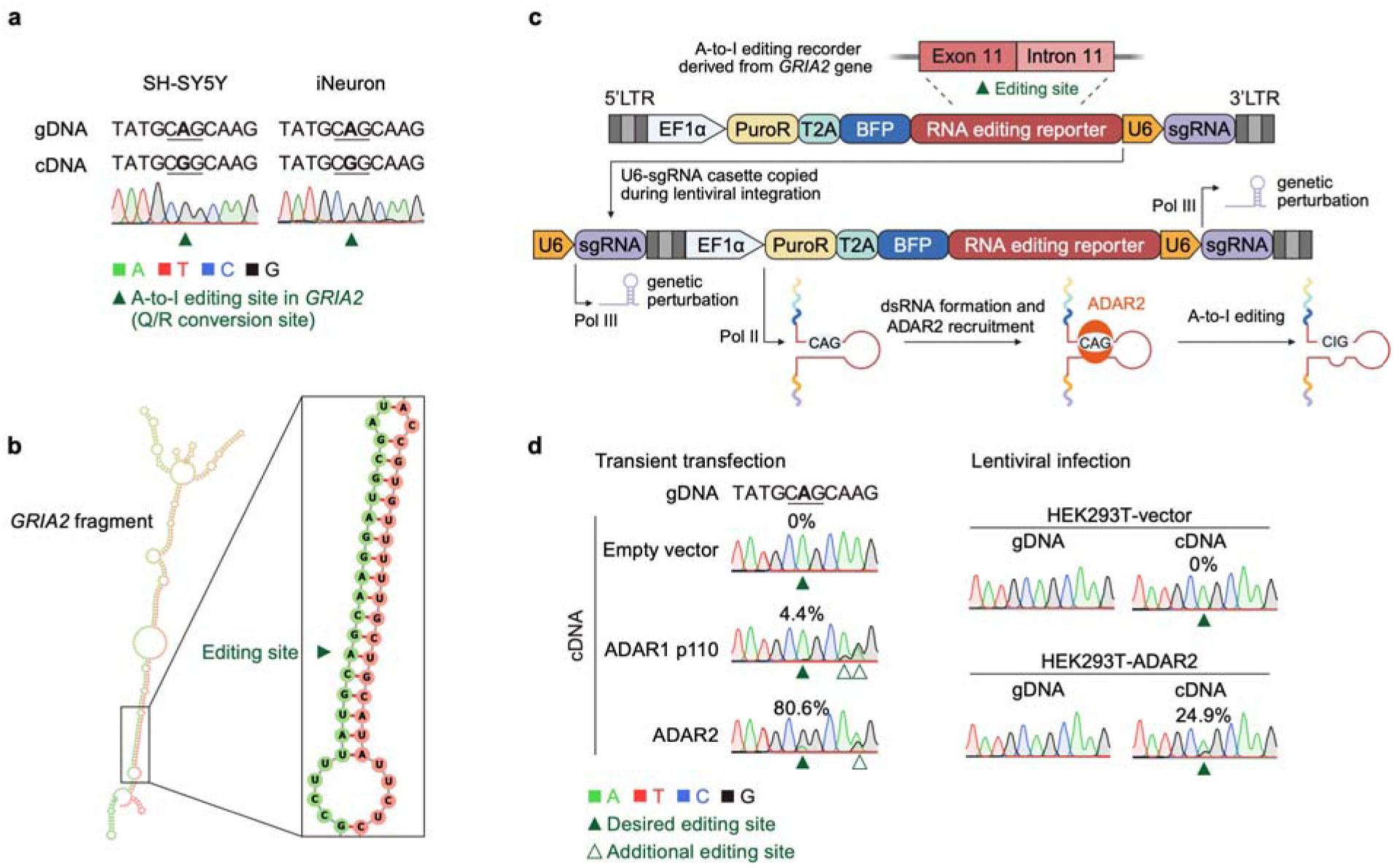
Development of CREDITS vector. **a**, Sanger sequencing electropherograms showing A-to-G conversion at the *GRIA2* Q/R site in SH-SY5Y cells and human iPSC-derived neurons (iNeurons). **b**, Predicted RNA secondary structure of the *GRIA2* fragment (chr4:157336674-157337074, hg38) using the *RNAfold* web server. **c**, Schematic of the CREDITS vector design. The *GRIA2*-derived A-to-I editing recorder is inserted into a CROP-seq vector (pMK1334), with EF1α promoter driving co-transcription of the recorder and sgRNA sequence. **d**, Sanger sequencing electropherograms showing A-to-G conversions in the CREDITS recorder in cells expressing ADAR1/2 via transient transfection (left) or lentiviral infection (right).

We inserted this A-to-I editing recorder into a previously published CROP-seq vector (pMK1334)^33^, resulting in the CREDITS vector in which the recorder can be co-transcribed with sequences containing sgRNA identity from a EF1a promoter, in addition to the U6 promoter-driven sgRNA expression for gene perturbation. Thereby, the effect of a specific genetic perturbation on A-to-I editing can be associating with the editing outcomes of the recorder (Fig. 1c). Importantly, the recorder region in the full-length CREDITS transcript is predicted to preserves the dsRNA structure (Supplementary Fig. 1a).

To test the performance of the CREDITS vector, we transiently transfected HEK293T cells with the CREDITS vector containing a non-targeting control sgRNA. Additionally, the cells were co-transfected with either an ADAR1 p110 or ADAR2 overexpression vector, or an empty vector. Following RNA extraction and reverse transcription, the recorder region was PCR amplified and subjected to Sanger sequencing. A strong A-to-I editing signal at the desired site in the recorder was detected in cells overexpressing ADAR2, whereas no or minimal editing can be detected in cells overexpressing either an empty vector or ADAR1 p110 (Fig. 1d). This is as expected because the Q/R conversion site in *GRIA2* is specific to ADAR2^15,32^, which is minimally expressed in HEK293T cells^34,35^. Interestingly, we observed additional editing signals at adjacent adenosines in cells overexpressing ADAR1 p110 and ADAR2, potentially due to high levels of transiently expressed ADAR proteins. To avoid ‘off-target’ editing, we sought to lower the level of ADAR2 overexpression by lentiviral delivering a ADAR2 expression cassette into the HEK293T cells, generating the HEK293T-ADAR2 cell line (Supplementary Fig. 1b-d). The CREDITS vector was also delivered by lentivirus into the HEK293T-ADAR2 cells (Supplementary Fig. 1e). As expected, a clear A-to-I editing signal was detected specifically at the desired site, with no ‘off-target’ editing sites detected (Fig. 1d). Moreover, no editing was detected in the genomic DNA of the recorder region, ruling out the possibility that the recorder was pre-edited during lentiviral production and infection prior to genome integration. Taken together, we established a sequencing-based reporter system that is suitable for pooled high-throughput CRISPR screening on A-to-I editing regulation.

### A CREDITS screen uncovered novel A-to-I RNA editing regulators

We next developed a HEK293T cell line that stably expresses ADAR2 along with the CRISPRi machinery (dCas9-BFP-KRAB), designated as CRISPRi-HEK293T-ADAR2. Using this cell line, we conducted a focused CRISPRi screen to identify factors that regulate A-to-I RNA editing. We constructed a CREDITS library containing 6,602 sgRNAs targeting 1,350 RNA-binding proteins (RBPs) and 250 non-targeting control sgRNAs. The CREDITS library was introduced into the CRISPRi-HEK293T-ADAR2 cells through lentiviral infection. Following selection and expansion, we extracted total RNA from the cells and performed reverse transcription. We then PCR-amplified the region containing both the RNA editing recorder and sgRNA sequence to generate an NGS library for paired-end sequencing (Fig. 2a). From each read pair, Read2 was used to identify the sgRNA identity, while Read1 to reveal the editing outcome of the recorder. The recorder editing level for each sgRNA was calculated as the ratio of edited to unedited read counts, and the phenotype of each sgRNA was determined as its relative editing level compared to non-targeting control sgRNAs (Fig. 2a; Methods). The strong correlation (r = 0.78) between phenotype scores from duplicate screens indicated the robustness and reproducibility of our screen (Fig. 2b).

**Fig. 2:**
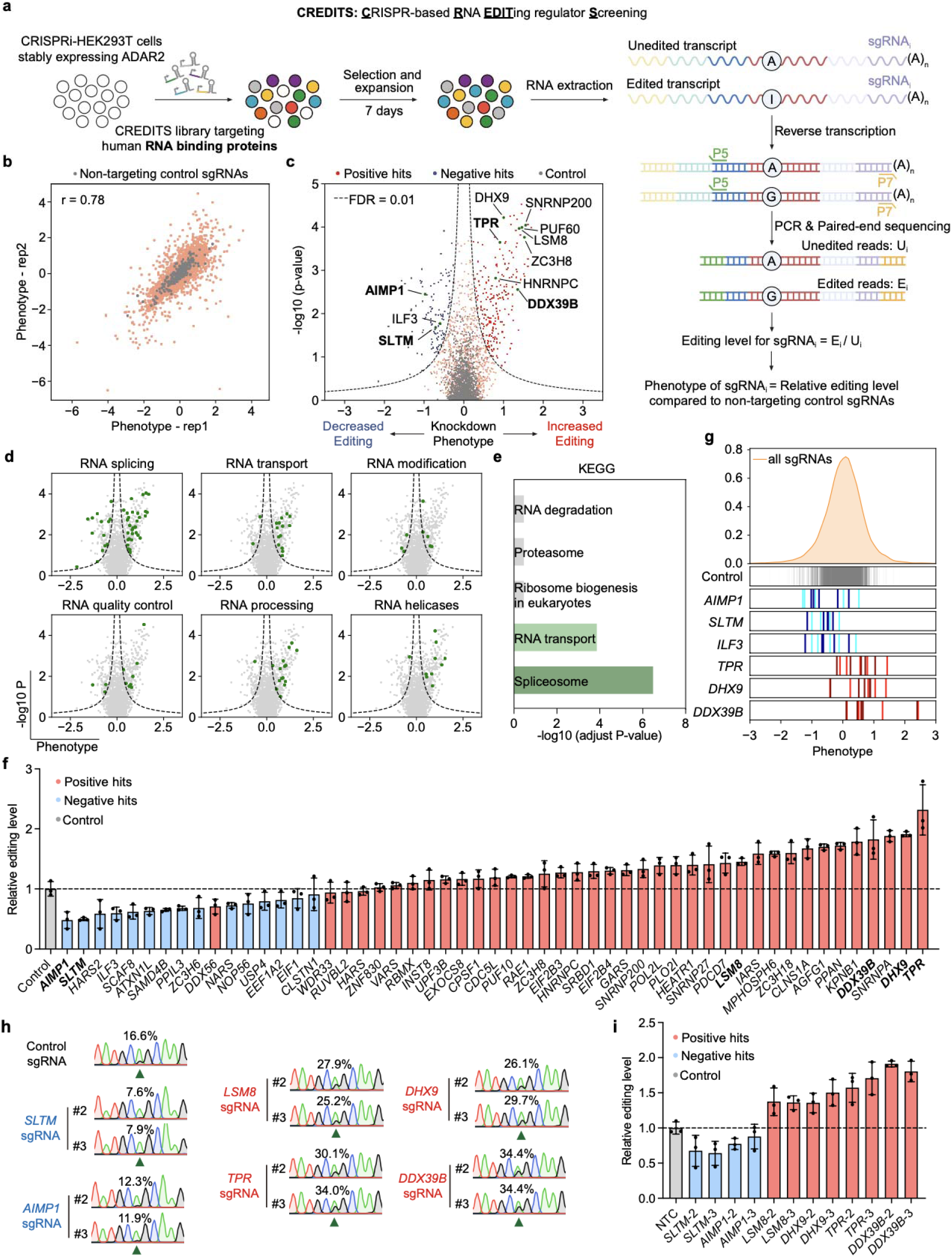
An RBP-focused CREDITS screen identifies known and novel RNA editing regulators. **a**, Screening strategy of CREDITS. HEK293T cells stably expressing ADAR2 and the CRISPRi machinery were transduced with an sgRNA library targeting 1,350 human RNA binding proteins (MOI < 0.3). Following puromycin selection and cell expansion, total RNA was extracted and reverse transcribed. A 1018 bp region containing both the editing recorder and sgRNA sequence was PCR-amplified to generate an NGS library for paired-end sequencing. For each sgRNA_i_, editing level was calculated as the ratio of edited reads (E_i_) to unedited reads (U_i_). The phenotype score for each sgRNA_i_ was determined by normalizing its editing level to the median editing level of non-targeting control sgRNAs. The screen was performed in two independent biological replicates. **b**, Scatter plot showing correlation of phenotype scores between two biological replicates (r = 0.78). **c**, Volcano plots showing knockdown phenotypes and statistical significance (Mann-Whitney U test) for genes targeted in the pooled screen. Screening results were analyzed by the MAGeCK-iNC pipeline. Dashed lines: cutoff for hit genes (FDR = 0.01). Known and novel hits discussed in the paper were labeled, and novel hits were highlighted in bold. **d**, Classification of hit genes based on RNA-related pathways. **e**, KEGG enrichment analysis for positive hits. Significant enriched signaling pathways were marked in green. **f**, Individual validation of hit genes from the primary screen using Sanger sequencing (mean ± s.d., n = 3 technical replicates). One sgRNA was cloned for each hit. Editing levels were normalized to that of the control sgRNA. Hit genes selected for secondary validation were highlighted in bold. **g**, Distributions of phenotype scores for non-targeting control sgRNAs and sgRNAs targeting selected negative (blue) and positive (red) hits. **h, i**, Validation of editing phenotypes for selected hits using Sanger sequencing with two additional sgRNAs per gene. (**h**) Representative electropherograms. (**i**) Quantification of editing levels relative to control sgRNAs (mean ± s.d., n = 3 technical replicates).

The screen identified 225 positive hits and 119 negative hits (FDR<0.01), whose knockdown increased or decreased A-to-I editing of the recorder, respectively (Fig. 2c; Supplementary table 1). These hits are involved in diverse RNA-related processes, including splicing, transport, modification, quality control, RNA processing and RNA helicase activity (Fig. 2d). Notably, our screen uncovered previously reported genes involved in A-to-I regulation, such as *ILF3*^28^, *PUF60*^31^, *HNRNPC*^31^, *SNRNP200*^31^, *ZC3H8*^31^, and *DHX9*^26^, validating the robustness of our screen. KEGG enrichment analysis revealed that positive hits were significantly enriched in Spliceosome and RNA transport pathways (Fig. 2e), in line with previous reports that inhibiting RNA splicing led to increased A-to-I editing^36,37^.

Intriguingly, the screen uncovered many hit genes that have not been previously associated with A-to-I editing. To validate these findings, we selected 53 hits (38 positive and 15 negative) for individual validation, encompassing both known and potentially novel A-to-I regulators. We individually cloned sgRNAs targeting these genes into the CREDITS vector and introduced them into CRISPRi-HEK293T-ADAR2 cells. Sanger sequencing analysis of the recorder’s editing outcomes showed remarkable consistency with the initial screen: all negative hits (15/15) and most of positive hits (34/38) reproduced their respective effects on A-to-I editing (Fig. 2f, g). The phenotypes of 6 selected genes that showed strong effect in both the initial screen and first-round validation-*SLTM*, *AIMP1*, *LSM8*, *TPR*, *DXH9,* and *DDX39B*-were further confirmed using two additional sgRNAs (Fig. 2h, i).

In summary, our screen uncovered known and potentially novel A-to-I RNA editing regulators, highlighting CREDITS as a robust platform for systematic investigation of RNA editing regulation.

### Development of scCREDIT-seq for scalable RNA editome-wide characterization of gene perturbations

Coupling CRISPR screening with single-cell omics enables screens on complex high-dimensional phenotypes. Technologies like CROP-seq and Perturb-seq, for instance, integrate CRISPR screening with single-cell RNA sequencing (scRNA-seq), allowing for transcriptome-wide characterization of gene perturbations at the single-cell level^38–41^.

Our initial screen relied on an exogenous reporter system. To examine whether and how hit genes modulate A-to-I editing of endogenous RNAs, we developed scCREDIT-seq (<u>s</u>ingle-<u>c</u>ell <u>CR</u>ISPR-based RNA <u>EDIT</u>ing <u>seq</u>uencing). This method leverages scRNA-seq to simultaneously profile sgRNA identity, gene expression signatures and A-to-I editing signatures at single-cell resolution. This approach enables high-throughput multiplexed characterization of transcriptome and editome changes across pooled CRISPR perturbations.

The workflow of scCREDIT-seq resembles that of CROP-seq (Fig. 3a). Briefly, sgRNAs are cloned into the CROP-seq vector pMK1334, which generates polyadenylated sgRNA-containing transcripts allowing for sgRNA detection via scRNA-seq. A pooled collection of cells expressing different sgRNAs is processed using a droplet-based capturing method, enabling the simultaneous capture of mRNA and polyadenylated sgRNA-containing transcripts from individual cells. During library construction, A-to-I editing events in the captured transcripts result in A-to-G conversions in the final library. The sgRNA-containing transcripts are additionally amplified and sequenced as previously described to facilitate sgRNA identity assignment^33,40,42,43^. Following sequencing, bioinformatic analyses are conducted to link sgRNA identity with gene expression and A-to-I editing profiles, and to determine transcriptome and editome changes associated with each perturbation.

**Fig. 3:**
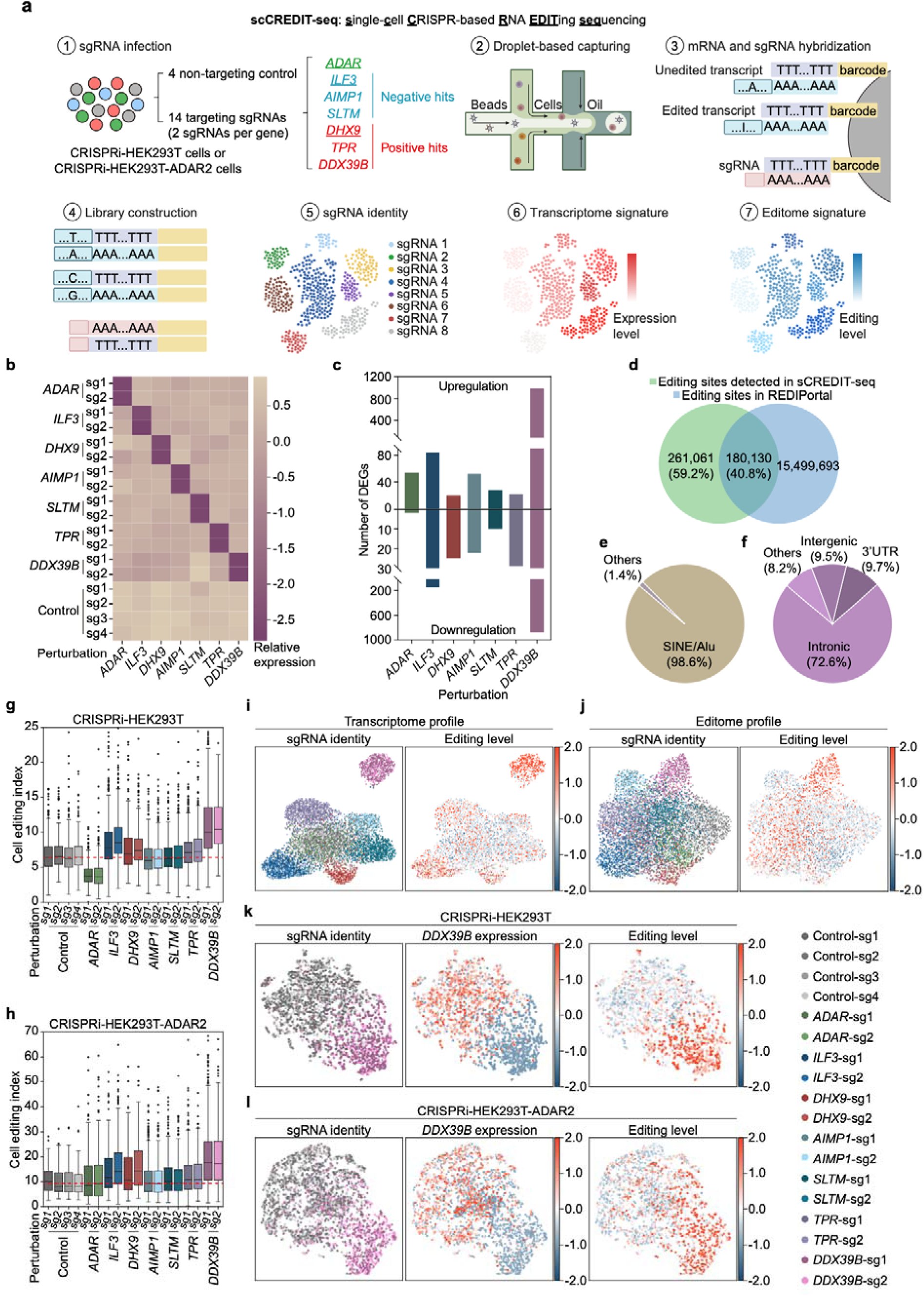
Single-cell transcriptome and editome characterization of A-to-I RNA editing regulators using scCREDIT-seq. **a**, Schematic of the scCREDIT-seq workflow. **b**, Heatmap showing on-target knockdown efficiency for each sgRNA in the scCREDIT-seq screen. **c**, Number of up- and down-regulated differentially expressed genes (DEGs) in each perturbation. **d,** Venn diagram showing the overlap between A-to-I editing sites identified by scCREDIT-seq analysis and known editing sites documented in the REDIportal database. **e, f**, Genomic distribution of high-confident RNA editing sites identified from the scCREDIT-seq. **g**, **h**, Box plots showing cell editing index (CEI) in cells expressing different sgRNAs in CRISPRI-HEK293T (up) and CRISPRI-HEK293T-ADAR2 (down) cells. The red dashed line represents the median CEI of the control cells. **i, j**, UMAP visualization of the scCREDIT-seq data following linear discriminant analysis (LDA) on transcriptome profile (**i**) or editome profile (**j**), color-coded by sgRNAs. The color legend representing each sgRNA was shown (bottom). **k**, **l**, UMAP visualization of a subset of the scCREDIT-seq data that contain only *DDX39B* sgRNAs and control sgRNAs following LDA in CRISPRI-HEK293T (**k**) and CRISPRI-HEK293T-ADAR2 (**l**) cells.

For our proof-of-principle scCREDIT-seq screen, we selected seven genes representing both known and novel regulators of A-to-I editing. As a positive control, we included *ADAR*, which encodes ADAR1, one of the primary enzymes responsible for catalyzing A-to-I editing. We selected another six genes from our initial screen: three positive hits (*DHX9*, *TPR*, and *DDX39B*) and three negative hits (*ILF3*, *AIMP1*, and *SLTM*). Among these, *ILF3*^28^ and *DHX9*^26^ had previously reported roles in A-to-I editing, while the other four genes (*TPR*, *DDX39B*, *AIMP1*, and *SLTM*) represent novel regulators identified through our screen. A total of 18 sgRNAs (two sgRNAs targeting each selected gene and four non-targeting control sgRNAs) were cloned into pMK1334, forming the scCREDIT-seq screen library. Parallel screens were conducted in HEK293T cells expressing the CRISPRi machinery, with or without ADAR2 overexpression (CRISPRi-HEK293T-ADAR2 and CRISPRi-HEK293T, respectively). Approximately 20,000 cells were processed for each scCREDIT-seq screen, and a total of 7,147 and 5,838 cells were retained after quality control for screens in CRISPRi-HEK293T and CRISPRi-HEK293T-ADAR2 cells, respectively. Since both screens yielded similar conclusions, we focused on the results from the CRISPRi-HEK293T cells, as they represent a more native context, unless otherwise specified.

We confirmed effective knockdown of all target genes in cells expressing the respective sgRNAs (Fig. 3b; Supplementary Fig. 2a). To determine gene expression changes induced by each perturbation, we performed pseudobulk differential gene expression analysis using DEseq2 with aggregated counts for each sgRNA. We found *DDX39B* knockdown induced the most dramatic transcriptome changes with 980 upregulated and 882 downregulated genes (Fig. 3c).

We developed a computational pipeline to identify A-to-I RNA editing sites from scRNA-seq data (Methods). Briefly, sequencing reads were aligned to the reference human genome, and A-to-G mismatches were identified from the bam files as potential A-to-I editing sites. These sites were then stringently filtered based on reads and cell representations and common genomics SNPs. Comparison with the REDIportal database^44^ revealed that of the 441,191 editing sites detected by scCREDIT-seq, 180,130 (41%) matched previously reported sites, while 261,061 (59%) were novel (Fig. 3d). We considered the overlapping sites as high-confident A-to-I editing sites and focused our subsequent analysis on these sites.

While most scRNA-seq methods use oligo-dT primers to target polyadenylated transcripts, these approaches can capture sequences beyond the 3’ end due to secondary priming positions throughout transcripts. Previous studies have shown that up to 25% of scRNA-seq reads may contain intronic or other gene body sequences^45^. Analysis of our scCREDIT-seq data confirmed this observation: while reads were enriched at the 3’ end, we detected substantial coverage across gene bodies and intergenic regions (Supplementary Fig. 3), suggesting that our approach can capture editing events throughout transcripts, despite the inherent bias of the sequencing method.

Analysis of genomic distribution of identified editing sites revealed that the vast majority (98.6%) of them were located within *Alu* repeats, a class of SINE (short interspersed elements) retroelements (Fig. 3e). These sites were predominantly found in intronic regions (72.6%), followed by 3’ untranslated regions (3’UTRs, 9.7%) (Fig. 3f). Similar genomic distribution of identified editing sites from scCREDIT-seq in CRISPRi-HEK293T-ADAR2 cells was observed as well (Supplementary Fig. 2b, c). This distribution pattern aligns with previous studies showing that *Alu* repeats, which are abundant in introns and 3’UTRs, serve as hot spots for A-to-I editing^46–48^, thus validating our approach. While most perturbations did not significantly affect the genomic distribution of editing sites, *ADAR* knockdown resulted in a slight increase in the proportion of editing events within 3’UTR regions (Supplementary Fig. 2d, e).

To evaluate how gene perturbations affect global RNA editing, we developed a cell editing index (CEI) to quantify overall editing levels in single cells (Methods). Analysis of CEI distributions showed that sgRNAs targeting the same gene produced similar effects, demonstrating high data reproducibility (Fig. 3g, h). As expected, *ADAR* knockdown markedly decreased CEI levels, while ADAR2 overexpression in CRISPRi-HEK293T-ADAR2 cells reversed this effect. Knockdown of all three positive hits—*DHX9*, *TPR*, and *DDX39B*—increased CEI levels, indicating their influence extends beyond the reporter to overall editing. Notably, *DDX39B* knockdown caused a dramatic increase in CEI levels in both CRISPRi-HEK293T and CRISPRi-HEK293T-ADAR2 cells, suggesting its strong regulatory role in global A-to-I editing.

Among the negative hits (*ILF3*, *AIMP1*, and *SLTM*), none showed decreased overall editing upon knockdown. Interestingly, knockdown of *ILF3* led to notable increases in global editing levels, contrasting with its effect on the reporter in our initial screen but consistent with its reported role as a negative regulator of RNA editing^28^. This finding suggests that ILF3 may exert site-specific effects on A-to-I editing, highlighting the value of scCREDIT-seq as a complementary approach to CREDITS for comprehensive editome characterization.

To map the landscapes of gene expression and A-to-I editing across different perturbations, we performed linear discriminant analysis (LDA), a supervised dimensionality reduction method that can maximize discrimination between different perturbations, followed by UMAP visualization. This analysis revealed distinct molecular signatures in both transcriptome and editome for each perturbation (Fig. 3i, j; Supplementary Fig. 2.f, g). Notably, *DDX39B* knockdown cells formed distinct clusters in both analyses, consistent with our findings that DDX39B knockdown induced the most pronounced overall transcriptome and editome changes among all perturbations. The same patterns persisted when the *DDX39B* perturbation was analyzed separately against control sgRNAs (Fig. 3k, l).

To determine the minimal cell coverage required for scCREDIT-seq, we performed a bootstrap analysis. We found that for perturbation with strong effects, such as *DDX39B* or *ADAR*, as few as 50 cells are sufficient to robustly distinguish them from control sgRNAs. In contrast, for perturbations with moderate effects, such as *ILF3*, more than 100 cells are needed (Supplementary Fig.S4).

In conclusion, we established scCREDIT-seq as a scalable and robust method for multiplexed characterization of transcriptome and editome changes for pooled CRISPR perturbations and identified *DDX39B* as a novel global repressor of A-to-I editing.

### Bulk RNA-seq confirms DDX39B as a global repressor of A-to-I RNA editing

To validate our scCREDIT-seq findings and further characterize how *DDX39B* knockdown affect A-to-I editing at different sites, we performed bulk RNA-seq on total RNAs from CRISPRi-HEK293T cells and CRISPRi-HEK293T-ADAR2 cells expressing either a control sgRNA or a *DDX39B-*targeting sgRNA. A-to-I editing sites were identified using the REDITools pipeline^49^, and only those also included in REDIportal were retained for downstream analysis.

As expected, A-to-I editing sites were predominantly found within *Alu* repeat regions (Fig. 4a), with the majority located in intronic regions (53.5%) and 3’UTRs (20.0%) (Fig. 4b). Among all mismatches identified, the frequency of A-to-G was significantly higher than those of other mismatches, such as A-to-T and A-to-C, which were negligible, confirming the detection of bona fide A-to-I events rather than technical artifacts. Knockdown of *DDX39B* significantly elevated overall A-to-I editing levels as measured by increased A-to-G conversions^50^, while A-to-T and A-to-C conversions remained unchanged (Fig. 4c), indicating a specific regulatory role of DDX39B on A-to-I editing, rather than a global effect on RNA or DNA fidelity.

**Fig. 4:**
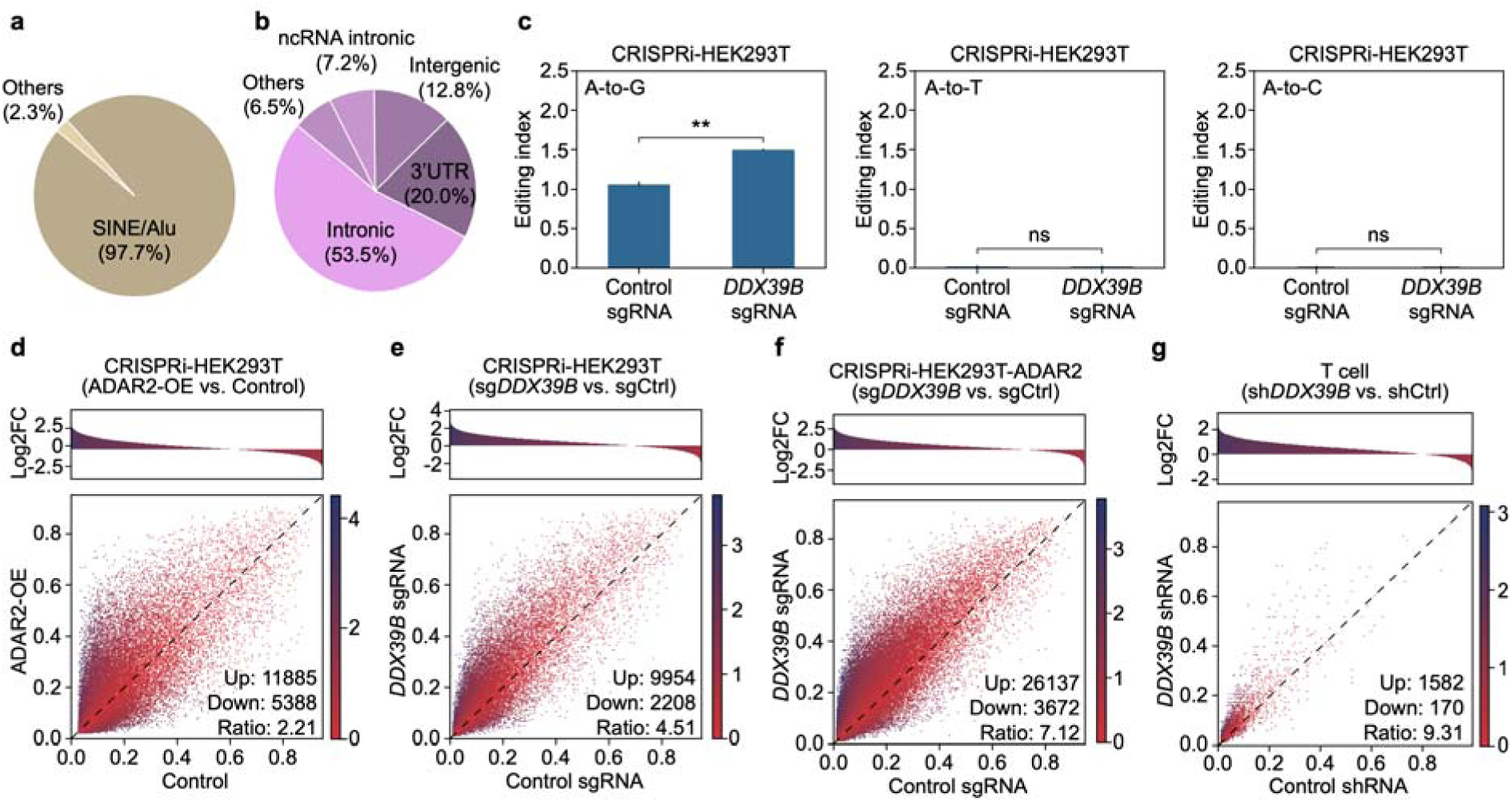
Bulk RNA-seq validation of DDX39B as a global repressor of RNA editing. **a**, **b,** Genomic distribution of A-to-I RNA editing sites detected from bulk RNA-seq. **c**, Histograms showing the A-to-G (left), A-to-T (middle), and A-to-C (right) conversion indices calculated by RNAEditingIndexer^50^. The A-to-G editing index is significantly higher in *DDX39B* knockdown cells compared to that of control cells. P value from Student’s t-test. **P < 0.01; ns, not significant. **d**, Differential RNA editing analysis between control and ADAR2-overexpressing (ADAR2-OE) cells at consensus editing sites. (**Top**) Histogram showing the distribution of log2 fold changes(log2FC) in editing levels (ADAR2-OE/control). (**Bottom**) Scatter plot comparing editing levels between control (x-axis) and ADAR2-OE cells (y-axis), where each dot represents an individual editing site. Colorbar representing log2FC was shown. Numbers of up-edited and down-edited sites (|log2FC| > 0.5) and their ratio were indicated. **e**-**g**, Differential RNA editing analysis for *DDX39B* knockdown in CRISPRi-HEK293T cells (**e**), CRISPRi-HEK293T-ADAR2 cells (**f**), and T cells (**g**). RNA-seq data for T cells were obtained from GSE145773^51^.

Next, we assessed A-to-I editing at the site-level. As expected, ADAR2 overexpression significantly increased the number of up-edited sites, yielding an up-edited to down-edited site ratio of 2.21 (Fig. 4d). Remarkably, *DDX39B* knockdown massively enhanced editing at over 80% of detected A-to-I sites, resulting in up-to-down ratios of 4.51 in CRISPRi-HEK293T cells and 7.12 in CRISPRi-HEK293T-ADAR2 cells (Fig. 4e, f). In contrast, the ratios between two biological replicates were close to 1 (Supplementary Fig. 5a, b). To validate the effect of DDX39B in other cell types, we analyzed published RNA-seq datasets for *DDX39B* knockdown in T cells (GSE145773^51^) and HeLa cells (GSE94730^52^). Consistently, *DDX39B* knockdown in these cells also significantly promoted global A-to-I editing (Fig. 4g; Supplementary Fig. 5c). These findings establish DDX39B as a global repressor of A-to-I RNA editing, validating our scCREDIT-seq findings.

### DDX39B interacts with ADAR1 in an RNA-dependent manner without altering ADAR1 expression or localization

To investigate how DDX39B regulates A-to-I editing, we first examined whether it affects the levels of ADAR proteins. Western blot analyses showed that neither *DDX39B* knockdown nor its overexpression affected ADAR1 levels in HEK293T cells (Fig 5a, b). Next, we investigated whether DDX39B physically interacts with ADAR1. Co-immunoprecipitation experiments revealed a strong association between DDX39B and ADAR1, which was diminished upon RNase A treatment (Fig. 5c), indicating that DDX39B interacts with ADAR1 in an RNA-dependent manner.

**Fig. 5:**
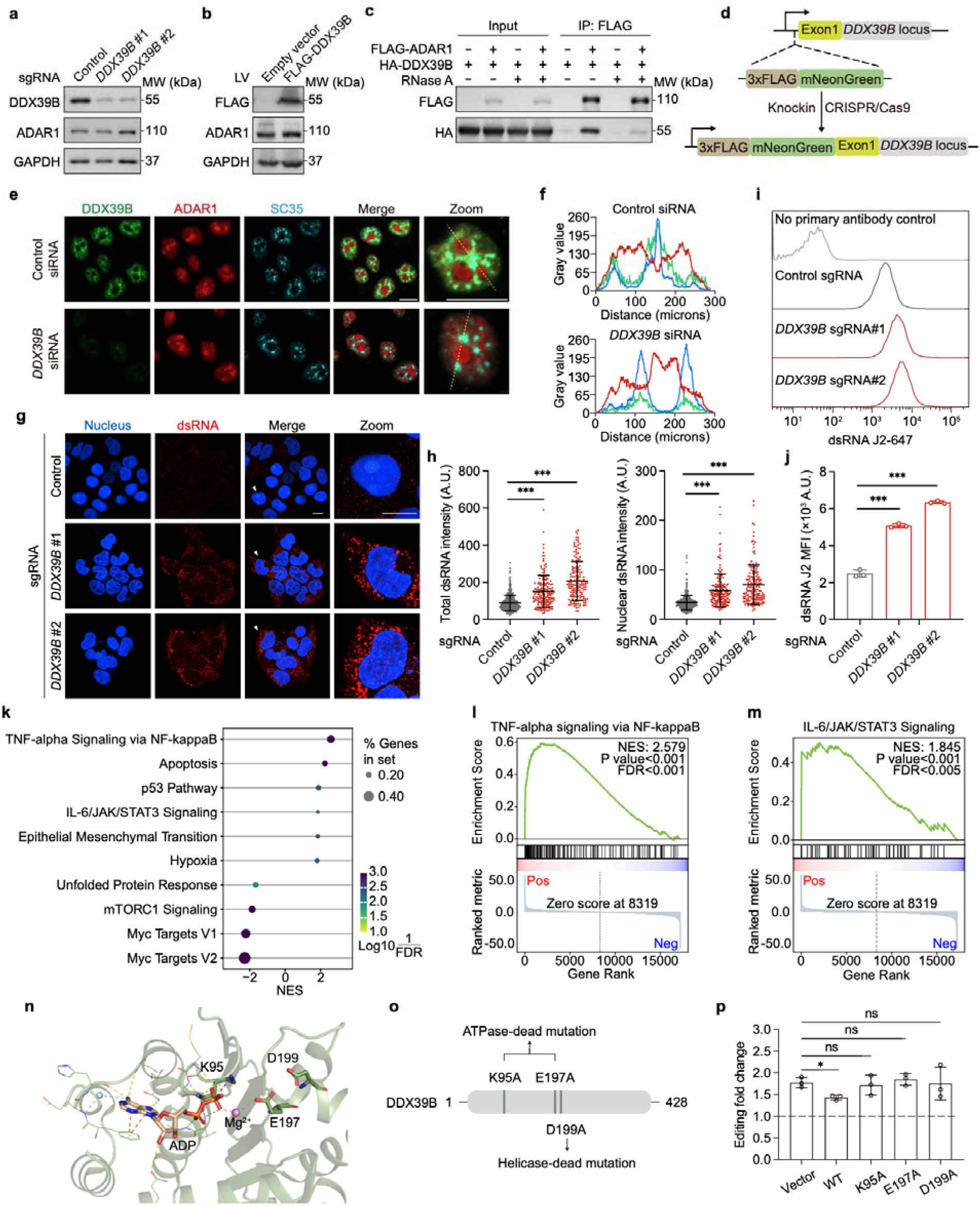
DDX39B regulates RNA editing through its helicase and ATPase activities. **a**, Western blot showing protein levels of DDX39B and ADAR1 in control and *DDX39B* knockdown HEK293T cells. GAPDH was used as the loading control. **b**, Western blot showing the levels of FLAG-DDX39B and ADAR1 proteins in CRISPRi-HEK293T cells with the overexpression of empty vector or FLAG-DDX39B. GAPDH was used as the loading control. **c**, Co-immunoprecipitation analysis showing the physical interaction between FLAG-ADAR1 (p110 isoform) and HA-DDX39B with or without RNase A treatment. **d**, Schematic of the generation of mNeonGreen-DDX39B HEK293T cells by CRISPR/Cas9-mediated knock-in. **e**, Immunofluorescence showing the localization of DDX39B (green), ADAR1 (red), and SC35 (cyan) in mNeonGreen-DDX39B HEK293T cells transfected with control or *DDX39B*-targeting siRNAs. **f**, Quantitative analysis of colocalization of DDX39B (green), ADAR1 (red), and SC35 (cyan) in mNeonGreen-DDX39B HEK293T cells transfected with control (top) or *DDX39B*-targeting siRNAs (bottom). **g**, Representative immunofluorescence images of dsRNA staining by the J2 antibody (red) in control and *DDX39B* knockdown HEK293T cells. Nuclei were counterstained with DAPI. Scale bar = 20 μm. **h**, Quantitative analysis of fluorescent intensity of dsRNA staining in whole cell (left) and in the nucleus (right). P values from unpaired two-sided Student’s t-test. ***P < 0.001. **i, j**, Flow cytometry analysis of intracellular dsRNA levels in control and *DDX39B* knockdown HEK293T cells. (**i**) Representative flow cytometry histograms showing dsRNA detection using J2 antibody staining. (**j**) Quantification of mean fluorescence intensity (MFI) from J2 staining (mean ± s.d., n = 3 biological replicates). P values from unpaired two-sided Student’s t-test. ***P < 0.001. **k-m**, Gene set enrichment analysis (GSEA) of DEGs in *DDX39B* knockdown cells (**k**). TNF-α signaling via NF-κB (FDR < 0.001, **l**) and IL-6/JAK/STAT3 signaling (FDR < 0.005, **m**) pathways were significantly upregulated. **n**, 3D structure of DDX39B highlighting the ATP-Mg^2+^-binding pocket (PDBid: 1xtj). **o**, Schematic of ATPase-dead and helicase-dead mutations in DDX39B. **p**, Rescue experiments showing RNA editing levels of the CREDITS reporter in *DDX39B* knockdown cells transfected with wild-type (WT) or enzymatic mutant DDX39B. Data were normalized to control cells and presented as fold change (mean ± s.d., n = 3 biological replicates). P values from unpaired two-sided Student’s t-test. *P < 0.05; ns, not significant.

Next, we examined the subcellular localization of DDX39B and ADARs. We generated a HEK293T cell line with its endogenous *DDX39B* locus tagged with a green fluorescence protein mNeonGreen at the N-terminus via CRISPR/Cas9-mediated homologous recombination (Fig. 5d). Immunofluorescence (IF) analyses in these cells showed that both DDX39B and ADAR1 were localized in the nucleus (Fig. 5e; Supplementary Fig. 6a, b;). Consistent with previous reports^53^, ADAR1 was distributed throughput the nucleus with a strong enrichment in the nucleolus. DDX39B, however, was localized in the nucleoplasm and enriched in nuclear speckles (stained by SC35), but excluded from the nucleolus, suggesting DDX39B may interact with ADAR1 in the nucleoplasm or nuclear speckle where they were co-localized. ADAR2 showed similar localization patterns as ADAR1 when overexpressed in HEK293T cells (Supplementary Fig. 6a, c). The localization and subnuclear distribution of ADAR1 remained unchanged upon *DDX39B* knockdown in HEK293T cells (Fig. 5e, f). Together, these findings excluded the possibility that DDX39B regulates A-to-I editing by modulating ADAR expression or localization.

### DDX39B regulates A-to-I editing through its helicase and ATPase activities

DDX39B is a member of the DEAD-box RNA helicase family, catalyzing dsRNA unwinding and promoting R-loop clearance ^54–58^. We hypothesized that knockdown of DDX39B may lead to dsRNA accumulation, thus providing more substrates for A-to-I editing. To test this, we measured the dsRNA levels in control and *DDX39B* knockdown cells using the J2 antibody, which specifically recognizes dsRNAs. Indeed, *DDX39B* knockdown dramatically increased dsRNA levels in the cell, as measured by IF and flow cytometry (Fig. 5g-j). The accumulation of dsRNAs in the cell will stimulate innate immune response^59,60^. In agreement with this, DEGs from the RNA-seq analyses of *DDX39B* knockdown showed a strong enrichment in the immunity and inflammation related pathways, including TNF-alpha signaling via NF-kB and IL-6/JAK/STAT3 signaling (Fig. 5k-m).

The RNA helicase activity of DDX39B is dependent on its ATPase activity^56,58,61^. Previous mutagenesis studies have identified K95 and E197 residues are essential for its ATPase activity, and D199 is essential for its helicase function^51,56,58^ (Fig. 5n, o). To dissect whether these enzymatic activities are required for the function of DDX39B in regulating A-to-I editing, we transduced wild-type (WT), ATPase-dead (K95A and E197A), and helicase-dead (D199A) mutants of DDX39B into CRISPRi-HEK293T-ADAR2 cells expressing a CREDITS vector with either a control sgRNA or a sgRNA targeting *DDX39*B (Supplementary Fig. 7a). We found that only WT DDX39B could partially revert the effect of increased A-to-I editing caused by *DDX39B* knockdown, whereas ATPase-dead or helicase-dead mutants failed to do so (Fig. 5p). Interestingly, overexpression of mutant DDX39B upregulated RNA editing levels in control cells, and this effect was further amplified by *DDX39B* suppression (Supplementary Fig. 7b, c), suggesting a dominant negative effect of DDX39B in regulating A-to-I editing. In summary, our findings suggest that DDX39B regulates A-to-I editing potentially by preventing dsRNA formation through its helicase and ATPase activities.

### Targeting DDX39B as a strategy for improving RNA editing-based tools

Having established DDX39B as a potent repressor of A-to-I editing, we explored its potential as a target to enhance the efficiency of existing RNA editing-based tools. CellREADR (Cell access through RNA sensing by Endogenous ADAR) is a cell monitoring and manipulation tool that couples RNA detection with the expression of effector proteins, relying on A-to-I editing^62^. To monitor the efficiency of CellREADR, we implemented a dual-fluorescence reporter system similar to that described in the original publication (Fig. 6a). The reporter consists of a UbC promoter-driven transcript containing mCherry, a sense-edit-switch RNA (sesRNA), and EGFP. While mCherry is constitutively expressed, EGFP expression requires sesRNA hybridization to its target RNA. This hybridization recruits ADAR proteins, which edit an in-frame UAG stop codon in the sesRNA to a UIG tryptophan codon, enabling downstream translation. Thus, the efficiency of CellREADR can be quantified by measuring the EGFP/mCherry ratio. We generated CellREADR vectors for five previously validated sense-edit-switch RNAs (sesRNAs), targeting human *EEF1A1* exon 3, *TP53* exon 3, *TP53* 5’UTR, *ACTB* CDS and *ACTB* exon 2. These vectors were transfected into CRISPRi-HEK293T cells expressing either a control or *DDX39B* targeting sgRNA. Intriguingly, knockdown of *DDX39B* substantially increased EGFP/mCherry ratios by up to 3.51-fold across all CellREADR constructs as measured by flow cytometry (Fig. 6b, c; Supplementary Fig.8). These findings indicate that targeting DDX39B could enhance CellREADR efficiency.

**Fig. 6:**
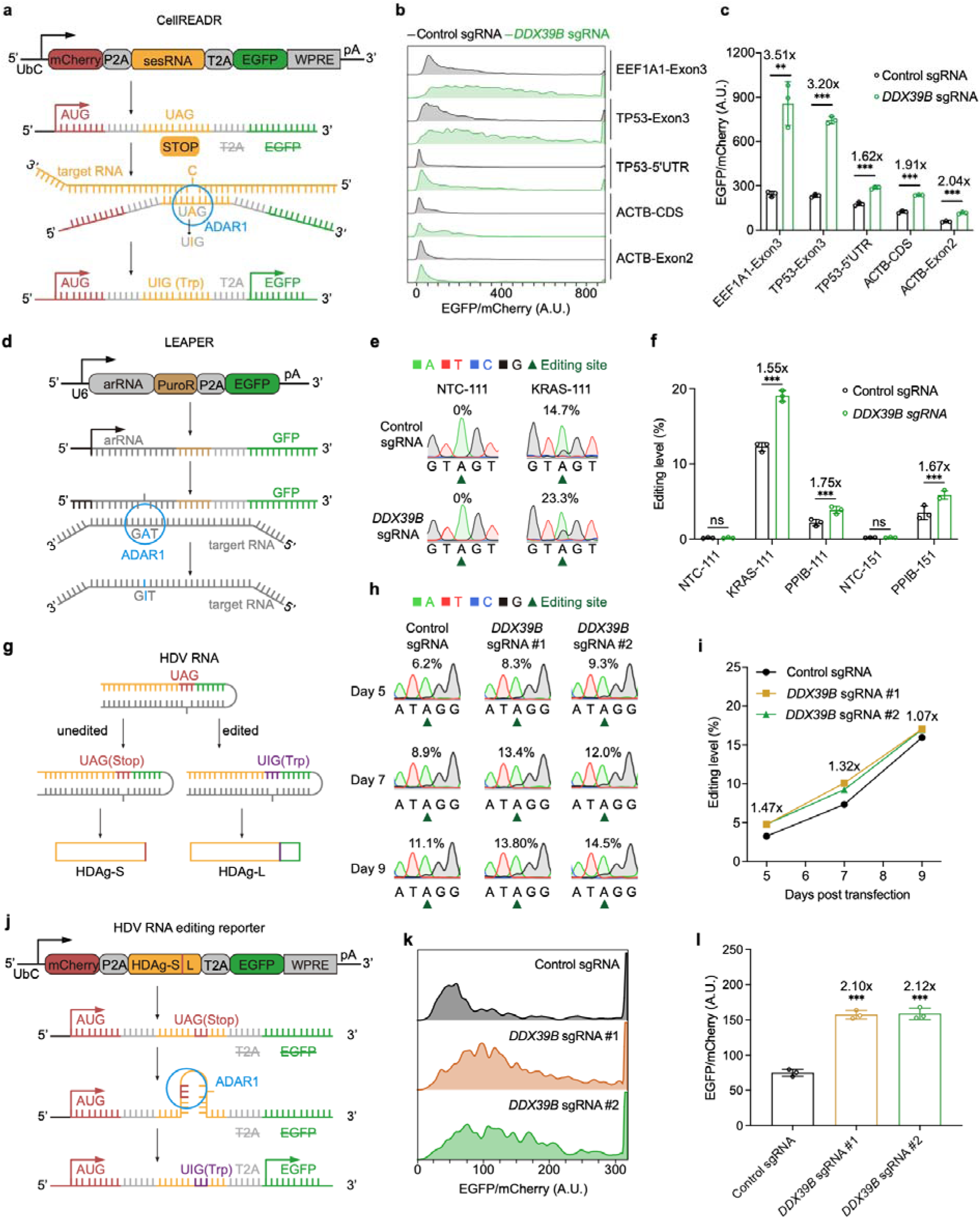
Targeting DDX39B as a strategy for improving RNA editing-based tools and developing anti-HDV therapy. **a**, Schematic showing the dual-fluorescence reporter for CellREADR. **b**, Flow cytometry analysis of CellREADR editing efficiency (EGFP/mCherry intensity ratios) in control and *DDX39B* knockdown cells. **c**, Quantification of CellREADR efficiency by flow cytometry (mean ± s.d., n = 3 biological replicates). P values from unpaired two-sided Student’s t-test. **P < 0.01; ***P < 0.001. Fold changes were indicated. **d**, Schematic illustration of LEAPER arRNA targeting an endogenous transcript. **e**, Sanger sequencing electropherograms showing A-to-G conversion at the LEAPER target site in control and *DDX39B* knockdown cells. **f**, NGS quantification of editing rates at targeted adenosines in *KRAS* and *PPIB* transcripts using arRNAs of varying lengths (mean ± s.d., n = 3 biological replicates). P values from unpaired two-sided Student’s t-test. ***P_<_0.001; ns, not significant. Fold changes were indicated. **g**, Schematic illustration of A-to-I editing in HDV RNA. ADAR1-mediated editing at the amber/W site extends HDAg-S to produce HDAg-L. **h**, Sanger sequencing electropherograms showing A-to-G conversion in HDV RNA from control and *DDX39B* knockdown cells. Target adenosine was highlighted. **i**, Time course analysis of HDV RNA editing levels in control and *DDX39B* knockdown cells by NGS. Average fold changes were indicated. **j**, Schematic of a dual-fluorescence reporter for measuring HDV genome A-to-I editing. The reporter replaces sesRNA in CellREADR with the HDV genome, triggering EGFP expression upon editing. **k, l**, Flow cytometry analysis of HDV reporter editing efficiency (EGFP/mCherry intensity ratios) in control and *DDX39B* knockdown HEK293T cells. (**k**) Representative flow cytometry histograms showing the distribution of EGFP/mCherry intensity ratio. (**l**) Quantification of HDV reporter editing efficiency in control and *DDX39B* knockdown HEK293T cells (mean ± s.d., n = 3 biological replicates). P values from unpaired two-sided Student’s t-test. ***P < 0.001. Fold changes were indicated.

LEAPER (leveraging endogenous ADAR for programmable editing of RNA) is another tool that employs a short, engineered ADAR-recruiting RNA (arRNA) to recruit endogenous ADAR proteins for targeted A-to-I editing^63,64^. We generated vectors expressing previously validated arRNAs targeting endogenous *KRAS* and *PPIB* transcripts (Fig. 6d) and transfected them into control and *DDX39B* knockdown CRISPRi-HEK293T cells. The transfected cells were sorted by FACS and the editing levels of the target RNAs were determined by Sanger sequencing and NGS. Notably, knocking down DDX39B significantly improved target RNA editing across all arRNAs tested (Fig. 6e, f), suggesting that targeting DDX39B could also enhance LEAPER efficiency.

Taken together, these results demonstrate the potential of targeting DDX39B as a strategy for improving RNA editing-based tools.

### Targeting DDX39B as a strategy for anti-HDV therapy

While A-to-I editing plays a crucial role in regulating host antiviral immunity, certain viruses have evolved to exploit the host’s A-to-I editing machinery to facilitate their life cycle. A notable example is the hepatitis D virus (HDV), which possesses a single-stranded circular RNA genome of approximately 1,700 nucleotides that folds into a unique rod-like structure through extensive base-pairing. The genome encodes only one protein, the HDV antigen (HDAg) which exists in two isoforms: HDAg-S (small) and HDAg-L (large), essential for viral replication and viral particle assembly, respectively. The switch from HDAg-S to HDAg-L relies on host ADAR1-mediated A-to-I RNA editing at a specific site known as the amber/W site^65,66^, resulting in a 19- to 20-amino acid extension at the C terminus (Fig. 6g). Maintaining a balanced level of A-to-I editing is critical for viral persistence, as HDAg-L, while necessary for viral particle secretion, strongly inhibits HDV replication. Therefore, modulating A-to-I editing represents a promising strategy for anti-HDV therapeutic development.

To investigate whether targeting DDX39B could influence A-to-I editing in the HDV genome, we transiently transfected a vector expressing the HDV RNA genome into control and *DDX39B* knockdown CRISPRi-HEK293T cells. At various time points post-transfection, we assessed the A-to-I editing levels at the amber/W site in the HDV genome using Sanger sequencing and NGS. In control cells, we detected A-to-I editing at the amber/W site in a time-dependent manner (Fig. 6h, i), resembling the phenomenon seen during live virus infection^67^, thus validating our approach. Intriguingly, knockdown of *DDX39B* markedly increased A-to-I editing levels in HDV RNA across all time points (Fig. 6h, i). This effect was further validated using a dual-fluorescence reporter system, analogous to the one used for CellREADR, but with the sesRNA sequence replaced by the HDAg sequence (Fig. 6j). In this reporter, EGFP expression requires A-to-I editing of an in-frame UAG stop codon in HDAg to UIG. Flow cytometry analysis revealed that *DDX39B* knockdown significantly increased the EGFP/mCherry ratio in CRISPRi-HEK293T cells expressing the reporter (Fig. 6k, l).

Collectively, these findings demonstrate that *DDX39B* suppression enhances A-to-I editing in the HDV RNA, suggesting its potential as a therapeutic target for anti-HDV therapy.

## Discussion

In this study, we developed two complementary CRISPR-based screening platforms to systematically identify and characterize key regulators of A-to-I RNA editing: CREDITS, which enables genome-scale screens using a reporter system, and scCREDITS-seq, which provides single-cell editome characterization for focused gene sets. Applying these platforms to screen 1,350 human RBPs, we uncovered multiple known and novel A-to-I regulators. Through detailed mechanistic investigation of one novel regulator, the RNA helicase DDX39B, we elucidated that it represses global A-to-I editing by preventing double-stranded RNA accumulation. We also demonstrated that targeting DDX39B has promising potential for enhancing RNA editing-based tools and for the development of therapies against HDV.

While the enzymatic mechanism of A-to-I editing by ADAR proteins is well characterized, the cellular mechanisms governing precise spatiotemporal control of editing events remain largely unknown. Previous efforts to identify RNA editing regulators have primarily employed three strategies: correlation analyses between genetic variants or gene expression and RNA editing levels^8,68^, editome analyses of publicly available or inhouse bulk RNA-seq data from various gene perturbations^31,69^, and proteomics and biochemical analyses of ADAR-interacting proteins^26–28^. Our platforms introduce the first genetic screening approach to studying A-to-I editing, offering a novel and systematic method to explore this regulatory landscape.

Our CREDITS method offers several key advantages. First, it enables NGS-based RNA-level readout by directly linking an RNA recorder to sgRNA, which is more straightforward and effective compared to fluorescence-based strategies that require converting RNA-level phenotypes into fluorescent signals and subsequent cell sorting for screening. This capability is particularly beneficial for cell types that are difficult to sort, such as neurons. Second, the method is highly versatile: by changing the recorder sequence, it can be easily adapted to study various DNA or RNA phenotypes. Indeed, similar screening approaches have successfully identified modulators of Prime editing^70,71^ and RNA m5C modification^72^.

The integration of single-cell technologies has transformed CRISPR screening by enabling complex high-dimensional phenotypes as readouts. Technologies such as Perturb-seq and CROP-seq focus on transcriptome responses, while the recently developed PerturbSci-Kinetics^41^ analyzes transcriptome kinetics and CPA-Perturb-seq^73^ examines polyA site usage. Our scCREDITS-seq method further expands the toolbox by providing a scalable method that can simultaneously assess transcriptome and editome changes in pooled genetic perturbations.

DDX39B is a multifaceted RNA helicase involved in multiple RNA-related processes, including RNA splicing and RNA export. Our study suggests that it prevents dsRNA accumulation through its helicase activity, thereby represses A-to-I editing. Further investigation is needed to elucidate how its functions in RNA splicing and export may contribute to this process. Furthermore, we demonstrated that DDX39B represents a promising target for improving RNA editing-based tools and developing anti-HDV therapies. Further studies are needed to develop DDX39B-targeting strategies, including Antisense Oligonucleotides (ASO), small molecular inhibitors, protein degraders, etc.

We anticipate that our screening platforms can be broadly applied to various cell types and conditions to decipher tissue-specific and context-dependent regulation of A-to-I editing, thereby advancing our understanding of RNA modification mechanisms, promoting the development of RNA modification-related technologies, and facilitating the development of therapeutic strategies for associated diseases.

## Methods

### Cell culture

HEK293T and SH-SY5Y cells were purchased from ATCC and cultured in Dulbecco’s Modified Eagle’s Medium and DMEM/F12 (Gibco) supplemented with 10% fetal bovine serum (TransGen) and 1% penicillin and streptomycin (Aladdin). iPSC was cultured and differentiated into neurons as described previously^33,43^. All cells were maintained at 37 ℃ and 5% CO_2_.

### Plasmid and siRNA transfection

The plasmids were transfected into the cells by Polyethylenimine Linear (PEI) MW40000 (Yeasen, 40816ES03) according to the standard protocol. The siRNAs targeting *DDX39B* (General biosystems) were transfected into the cells in combination with Lipofectamine RNAiMax Reagent (Invitrogen, 13778150) and Opti-MEM (Gibco, 31985-062) medium at 20 nM as final working concentration. The targeting sequences of each siRNA were listed in Supplementary Table 2.

### Generation of CREDITS vector

Sequence flanking the Q/R conversion site of *GRIA2* (chr4:157336674-157337074, hg38) was amplified from HEK293T genomic DNA by PCR. The amplified product was instead into a CROP-seq vector pMK1334 (Addgene, 127965) by replacing original WPRE cassette between EcoRI and SalI sites using ClonExpress® Ultra One Step Cloning Kit (Vazyme, C115).

### Generation of CRISPRi-HEK293T cell line with the stable overexpression of ADAR2

CRISPRi-HEK293T cell line was generated as described previously^74^. The cDNA of human *ADARB1* was gifted from Dr. Hao Chen (SUSTech) and was subcloned together with a 3×Flag tag into a pLV2-UBC-mCherry-Hyg-CMV-MCS vector (Miaoling, P36518) between BamHI and AgeI sites. The lentivirus was packaged using psPAX2 (Addgene, 12260) and pMD2.G (Addgene, #12259) plasmids as described previously^75^. 7 days post lentivirus transduction, mCherry-positive CRISPRi-HEK293T cells with ADAR2 overexpression was sorted by FACS (BD FACSAria SORP). The cell pool was cultured and enlarged and the ADAR2 overexpression efficiency was verified by qRT-PCR and WB. The CRISPRi-HEK293T-ADAR2 cells was integrated with CREDITS vector using lentivirus transduction for further experiments.

### Detection of RNA editing in CREDITS vector by sanger sequencing

Total RNA from the cells with CREDITS was extracted by MolPure^®^ Cell RNA Kit (Yeasen, 19231ES50) and 1 μg RNA was reverse transcribed to cDNA using HiScript III RT SuperMix (Vazyme, R323) according to the manufacturer’s instruction. To measure RNA editing level in CREDITS vector, the cDNA products were used as templates for PCR to amplify the fragments flanking the A-to-I editing site and the PCR products were subjected to sanger sequencing. The abundance of nucleotide A and G in sequencing results was determined in SnapGene software and the RNA editing level was defined as the percentage of G/(A+G).

### CRISPRi screening with CREDITS and data analysis

To integrate CREDITS vector with a human RNA binding protein (RBP) sgRNA library, 6853 unique sgRNA sequences targeting 1,350 RBPs along with 250 non-targeting control sgRNAs, was synthesized by GENEWIZ and cloned into CREDITS vector between BstXI and BlpI sites. To evaluate the library quality, the fragment harboring sgRNA sequence was amplified using Phanta Flash Master Mix (Vazyme, P520) as manufacturer’s instructions, and the PCR products were processed by next-generation sequencing (NGS).

For lentivirus production of the CREDITS vector integrated with RBP library, 5×10^6^ HEK293T cells were plated onto a 15cm dish for 24 hours before transfection. 15 μg library plasmid and 15 μg PackageMix plasmid were transfected into the HEK293T cells using Polyethylenimine Linear (PEI) MW40000 (Yeasen, 40816ES03). The PackageMix was prepared as an equal ratio mixture of three plasmids, including pMDLg/pRRE (Addgene, 12251), pRSV-Rev (Addgene, 12253), and pMD2.G (Addgene, 12259). 48 hours later, the supernatant containing lentivirus was collected and filtered with a 0.45 μm filter (Millipore, SLHV033RB).

The lentivirus was transduced into CRISPRi-HEK293T-ADAR2 cells at 0.3 multiplicity of infection (MOI). 48 hours later, the transduced cells were selected with 2 µg/mL of puromycin for 48 h to eliminate uninfected cells and generate a genome-edited cell pool. After 7-day passage in medium containing no puromycin, 10 million cells were collected for total RNA extraction by TRIzol reagent (Ambion). The mRNA containing poly-A tail from a total of 20 μg extracted RNA was reversed transcribed into cDNA using Oligo-dT primers by TransScript^®^ II One-Step gDNA Removal and cDNA Synthesis SuperMix (TransGen, AH311). All synthesized cDNA was used as template for PCR to amplify the linear region containing RNA editing reporter and sgRNA sequence. The PCR products were prepared for paired-end sequencing with Illumina platform (BerryGenomics). Raw FASTQ files for Read2 were cropped and aligned to the sgRNA reference of the RBP library using bowtie (v1.1.2) to identify sgRNAs, while Read1 files were cropped and aligned to a reference containing edited and unedited A-to-I reporter sequences to determine editing outcomes. This analysis enabled the determination of reporter editing levels for each identified sgRNA. Subsequently, the MAGeCK-iNC pipeline^33^ was implemented to assess sgRNA-level and gene-level editing phenotypes as compared to non-targeting controls.

### Primary validation of screening hits

Individual sgRNAs of top negative and positive hits in analysis results were cloned into the CREDITS vector via BstXI and BlpI sites. For lentivirus production of the CREDITS vector integrated with individual sgRNA, 1×10^5^ HEK293T cells per well were plated onto a 12-well plate for 24 hours before transfection. 0.5 μg CREDITS vector and 0.5 μg PackageMix plasmid were transfected into the HEK293T cells using PEI. The remaining procedures were performed as described above. 5-day post lentivirus transduction, the cells were collected for RNA extraction and the RNA editing level was determined as described in “Detection of RNA editing in CREDITS vector by sanger sequencing”. The sgRNA sequences used in this study was listed in Supplementary Table 2.

### Quantitative real**lZI**time polymerase chain reaction (qRT-PCR)

Total cellular RNA from cells was extracted using MolPure® Cell RNA Kit (Yeasen, 19231ES50) following the manufacturer’s instructions. cDNA was reverse transcribed from 1 μg RNA using HiScript III RT SuperMix for qPCR (Vazyme, R323). Quantitative real-time PCR was performed using Taq Pro Universal SYBR qPCR Master Mix (Vazyme, Q712) according to the manufacturer’s protocol and ran on the LineGene 9600 Plus Real-Time PCR Detection System (Bioer, FQD-96A). Gene relative expression level was calculated using the 2^−ΔΔCt^ method, and *ACTB* was used as an endogenous control. The qRT-PCR primers used in this study are listed in Supplementary Table 4.

### Immunoblotting

Cells were lysed in RIPA buffer (Beyotime, #P0013B) supplemented with phosphatase inhibitor cocktail (Selleck, B15001) and protease inhibitor cocktail (MCE, HY-K0010). The protein concentration of each sample was determined by Easy II Protein Quantitative Kit (TransGen, DQ111). Protein was denatured in loading buffer (Solarbio, P1041) by boiling at 95 ℃ for 10 minutes, and 20 μg protein from each lysate was electrophoresed on SDS-PAGE gels and was transferred to 0.45 µm PVDF membranes (Immobilon, IPVH00010). The membrane was blocked by 5% BSA (Sigma-Aldrich, V900933) in TBST (Sangon, C520009), and incubated with primary antibodies as indicated at 4 ℃ overnight. Next day, the membrane was incubated with horseradish peroxidase–conjugated secondary antibodies for 1 hour at room temperature. The protein signal was detected using the clarity western ECL substrate (EpiZyme, SQ202L). The antibodies used in this study are summarized here: anti-ADAR1 (CST, 81284S), anti-Flag (Absin, abs830014), anti-HA (CST, 3724S).

### Co-immunoprecipitation (Co-IP)

The experiment was performed as described previously^75^. In brief, 3×10^6^ HEK293T cells were plated onto a 10cm diameter dish for 24 hours before transfection. The pcDNA3.1-3×Flag-ADAR1 and pcDNA3.1-3×HA-DDX39B were transfected into the cells. 48 hours later, the cells were collected and lysed in NP-40 buffer (Beyotime, P0013F). The cell lysates were centrifuged at 12,000 rpm at 4 ℃ for 10 minutes, then the supernatant was collected and treated with 400 μg RNase A (Yeasen, 10406ES03) at room temperature for 2 minutes. Subsequently, the supernatant was incubated with anti-Flag magnetic beads (Beyontime, P2181S) at 4 ℃ for 4 hours. After washing, the beads affiliated with proteins were denatured and subjected to immunoblotting analysis.

### scCREDIT-seq and data analysis

The overall scCREDIT-seq process was derived from CROP-seq as described previously^33^. In brief, CRISPRi-HEK293T cells with or without ADAR2 overexpression were transduced with individual lentivirus of selected sgRNAs in the CROP-seq vector pMK1334. After puromycin selection and expansion, all cells were pooled together at equal ratio. Approximately 20,000 cells were loaded into microfluidic chip of Chip A Single Cell Kit v2.1 (MobiDrop, S050100301) to generate droplets with MobiNova-100 (MobiDrop, A1A40001). Each cell was involved into a droplet which contained a gel bead linked with up to millions oligos (cell unique barcode). After encapsulation, droplets suffer light cut by MobiNovaSP-100 (MobiDrop, A2A40001) while oligos diffuse into reaction mix. The mRNAs were captured by cell barcodes with cDNA amplification in droplets. Following reverse transcription, cDNAs with barcodes were amplified, and a library was constructed using the High Throughput Single-Cell 3’ Transcriptome Kit v2.1 (MobiDrop, S050200301) and the 3’ Dual Index Kit (MobiDrop, S050300301).

To facilitate sgRNA assignment, sgRNA-containing transcripts were additionally amplified by hemi-nested PCR reactions as described previously^33^. The sgRNA-enrichment libraries were separately indexed and sequenced as spike-ins alongside the whole-transcriptome scRNA-seq libraries using NovaSeq 6000 system. The primers used for sgRNA enrichment PCR was listed in Supplementary Table 3.

#### Processing of single-cell gene expression data

Single-cell RNA sequencing (scRNA-seq) and Feature Barcoding data from HEK293T cells were processed using STARsolo (STAR v2.7.11a)^76^. Human genome GRCh38 and GENCODE v44 were used as references^77^. The gRNA-amplified library data was analyzed and assigned to cell barcodes using customized Python3 scripts (available at https://github.com/comics-bio/scCREDITS-seq). The resulting count matrices and gRNA-cell barcode mapping matrices were then used as input for downstream analyses with the scanpy package (version 1.10.2)^78^. We retained cells with unique sgRNA assignments and filtered out those that were not fully perturbed. To further refine our dataset, we excluded cells that may have received an sgRNA but did not exhibit strong molecular evidence of successful perturbation. Specifically, we retained only cells where the expression levels of target genes were below those of 75% of cells. Conversely, the cells with control sgRNA were required to have all target genes expressed in more than 85% of cells.

For visualization, we used Linear Discriminant Analysis (LDA), a supervised dimensionality reduction technique that identifies a low-dimensional subspace to maximize discrimination between different groups (‘perturbations’) in the data. Before applying LDA, we reduced the dimensionality of our dataset (editing index or expression matrix) to 108 features (6 projected principal components × 18 perturbations). The components returned from LDA were then used as input for 2D visualization with UMAP.

#### Single-Cell Metric for A-to-I RNA Editing

We developed a single-cell metric for A-to-I RNA editing that adjusts for the total gene abundance to prioritize the detection of overall changes in RNA editing. We used only reads with a mapping quality greater than 255 and a base quality above 30 as input. Our analysis focused on known editing sites reported in REDIportal^79^ and we excluded common A to G SNPs annotated in dbSNP (version v156)^80^. Highly confident A-to-I sites were retained through a series of filtering steps. Specifically, we only kept sites that met all of the following criteria: (i) >=2 edited reads across all cells; (ii) >=10 reads (edited and unedited) across all cells; (iii) editing detected in >=0.5% of all cells; (iv) at the editing site, the number of cells with base C/T (noise) < 20% * the number of cells with base G (signal). The remaining sites were annotated with gene symbols using RefGene, repeat regions using RepeatMasker^81^, and known RNA editing sites using the REDIportal (V2)^79^ via ANNOVAR (version 2020.06.07)^82^. The cell editing index (CEI) is defined as below:

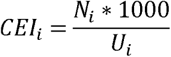

where:

*CEI_i_*: Cell Editing Index for cell i.

*N_i_*: Number of UMIs with at least one editing site in cell i.

*U_i_*: Total UMI counts of cell i.

Z-score normalized CEI values were used for visualization.

Statistical testing for CEI between different sgRNA groups was performed using a t-test, and Differential Expression Gene (DEG) analysis was conducted with the decoupleR (Version 1.8.0)^83^ and PyDESeqE (Version 0.4.12)^84^ with a pseudobulk stragecy.

### Bulk RNA sequencing (RNA-seq) and data analysis

The CRISPRi-HEK293T cells with different gene perturbation by sgRNAs were collected and total RNA was extracted using TRIzol reagent (Ambion). The sgRNA sequences used in this experiment were listed in Supplementary Table 2. RNA quantification and concentration was measured using Qubit 4.0 (Thermo Scientific). RNA integrity was assessed using Agilent 2100 Bioanalyzer (Agilent Technologies). RNA purity was controlled by NanoDrop.

A total amount of 500 ng RNA per sample was used as input material for the RNA sample preparations. Ribosomal RNA was removed from total RNA using Ribo-off rRNA Depletion Kit (Yeasen). Sequencing libraries were generated using Hieff NGS^®^ Ultima Dual-mode mRNA Library Prep Kit for Illumina^®^ (Yeasen, 12301) following manufacturer’s recommendations and index codes were added to attribute sequences to each sample. The library quality was assessed on the Qseq 100 system. The dsDNA library was denatured, cyclized, and digested to obtain single-stranded circular DNA, and DNB nanospheres were obtained by Rolling Circle Amplification (RCA). The prepared DNB was loaded onto a microarray chip (Patterned Array) and sequenced on DNBSEQ-T7 platform (Geneplus-Shenzhen) using Combinatorial Probe-Anchor Synthesis (cPAS), and 150 bp paired-end reads were generated.

All raw data were mapped to the GRCh38 genome assembly using GENCODE gene annotations v44 with STAR (v2.7.11a). The resulting BAM files for each sample were sorted with samtools^85^ and used as input for RNAEditingIndexer (latest version)^50^ to assess sample RNA editing level. A-to-I RNA editing events and editing levels at known sites with coverage above ten reads were detected and queried using REDITools^49^. To minimize false positives, only editing sites included in REDIportal^79^ were used for downstream analysis. Subsequently, all selected sites were annotated with ANNOVAR^82^ to map gene symbols using RefGene and repeat regions using RepeatMasker^81^ data. All downstream analyses were performed using customized Python3 scripts (https://github.com/comics-bio/scCREDITS-seq).The numbers of up- and down-regulated editing sites were calculated only for sites with an absolute log2 fold change (|log2FC|) greater than 0.5. For gene expression analysis, gene quantification was performed with RSEM (v1.3.1). The R package DESeq2 was then used to fit the expression count matrix to a negative binomial distribution and identify differentially expressed genes. Adjusted p < 0.05 and log2 fold-change > 1 were used to determine statistical significance. Gene Set Enrichment Analysis (GSEA) was done with GSEAPY (version 1.1.3).

### mNeonGreen knock-in for DDX39B in HEK293T

The CRISPR/Cas9 system was used to generate engineer HEK293T cell line. sgRNA targeting the exon 1 of *DDX39B* was cloned into PX459 (Addgene, #62988). Donor double-strand DNA (dsDNA) containing microhomology and mNeonGreen cassette was constructed. To generate mNeonGreen-*DDX39B* endogenous tagged HEK293T, 2ug PX459 and donor dsDNA were transfected into HEK293T using PEI (Yeasen,40816ES01). 48hrs later, HEK293T cells were treated with 2 ug/ml puromycin for 72hrs. Green-fluorescent single cells were sorted into a 96-well plate by FACS (BD FACSAria SORP). One week later, homogeneous knock-in clones were verified by PCR and Sanger sequencing. The genotyping PCR primers were listed in Supplementary Table 4.

### Immunofluorescence (IF)

Cells were plated at 5×10^4^ per well on sterilized, Matrigel-coated 12mm diameter round glass coverslips (CITOTEST, 10210012CE) in 24-well plates. Next day, cells were fixed with 4% paraformaldehyde (Beyotime, P0099) at room temperature for 15 minutes. After washing, cells were treated with 0.1% Triton X-100 (Coolaber, CT11451) in PBS (BBI, E607008) for 5 minutes, then the cells were blocked with 3% BSA (Sigma-Aldrich, V900933) in PBS (BBI, E607008) for 30 minutes. The cells were incubated with primary antibodies as indicated at 4℃ overnight. Next day, the primary antibodies were washed out and the secondary antibodies were applied to the cells at room temperature for 1 hour. After washing, the nucleus was stained by DAPI (Beyotime, C1006) at room temperature for 10 minutes. Finally, coverslips were then washed three times with PBS for 5 minutes each. One drop of Mounting Medium (SouthernBiotech, 0100) was added to the coverslips and the coverslips were sealed with slides (CITOTEST, 10127105P). The antibodies used in this study were listed here: anti-ADAR1 (CST, 81284S), anti-ADAR2 (SANTA CRUZ BIOTECHNOLOGY, sc-73409), anti-SC35 (Abcam, ab11826). The images were captured in a confocal microscope (Zeiss, LSM 980), and the protein colocalization analysis was performed by Fiji.

### Detection of dsRNA accumulation

To detect the dsRNA accumulation by IF, cells were treated as described above in “Immunofluorescence (IF)”. The J2 antibody (Merck, MABE1134) specifically recognizes dsRNA was used in this experiment. The images were captured in a confocal microscope (Nikon) and subjected to the deconvolution processing using default setting. The fluorescent intensity of dsRNA was quantified using CellProfiler.

To detect the dsRNA accumulation by flow cytometry, cells were detached with 0.5 mM EDTA (LABSELECT, BL518A) and then washed twice using PBS. Cells were fixed with 4% paraformaldehyde (Beyotime, P0099) at room temperature for 20 minutes. After washing, cells were permeabilized with 0.1% Triton-X-100 (Coolaber, CT11451) in PBS for 15 minutes followed by incubation in 3% BSA in PBS for 30 minutes, then the cells were stained with J2 antibody (SCICONS, 10010200) for 1 hour at ice and secondary antibody for 30 minutes at room temperature. The dsRNA intensity was measured using flow cytometry (BD FACSCanto SORP).

### Protein structure analysis and site mutagenesis

The crystal structure of DDX39B complexed with ADP and Magnesium ion was downloaded from PDB dataset (PDBid: 1xtj) and was analyzed using The PyMOL Molecular Graphics System Version 3.0 (Schrödinger). The site mutations of K95A, D197A, and E199A were introduced to wild-type DDX39B plasmid by homogenous recombination. The primers were designed as listed in Supplementary Table 4.

### Generation of CellREADR vector and data analysis

The CellREADR vector was generated by adapting a previously published article^62^. Briefly, the sequence of sense–edit–switch RNA (sesRNA) was synthesized by Tsingke and was inserted into FUW vector (Addgene, 14882) with mCherry and EGFP fragments using ClonExpress^®^ Ultra One Step Cloning Kit (Vazyme, C115). The sesRNA sequences and primers used in this study were listed in Supplementary Table 3 and 4, respectively.

The CellREADR vector was transfected into the cells using PEI. 72 hours later, the cells were collected for flow cytometry analysis (BD FACSCanto SORP). The cell conversion ratio was defined as the percentage of mCherry^+^/EGFP^+^ double-positive cells over mCherry^+^ single-positive cells.

### Generation of LEAPER vector and data analysis

The LEAPER vector was generated by adapting a previously published article^63^. In brief, the ADAR-recruiting RNAs (arRNA) was synthesized by Tsingke and was inserted into a pU6-pegRNA-GG-acceptor (Addgene, 132777) by replacing original mRFP1 cassette between dual BsaI sites. The hPGK promoter, puromycin resistance, EGFP, and SV40 poly(A) signal terminator were inserted to the vector using ClonExpress^®^ Ultra One Step Cloning Kit (Vazyme, C115).

The LEAPER vector was transfected into the cells using PEI. 72 hours later, the EGFP positive cells were sorted by FACS (BD FACSAria SORP), and total RNA was extracted using MolPure^®^ Cell RNA Kit (Yeasen, 19231ES50). 500 ng RNA was reversed transcribed into cDNA using cDNA using Oligo-dT primers by TransScript^®^ II One-Step gDNA Removal and cDNA Synthesis SuperMix (TransGen, AH311). The cDNA products were used as templates for PCR to amplify the fragments flanking the A-to-I editing site and the PCR products were subjected to sanger sequencing. The abundance of nucleotide A and G in sequencing results was determined in SnapGene software and the RNA editing level was defined as the percentage of G/(A+G). The arRNA sequences and primers used in this study were listed in Supplementary Table 3 and 4, respectively.

### HDV RNA editing detection

The HDV infectious clone was generated by inserting a 1.1mer HDV genome into the pcDNA3.1 plasmid. (kindly provided by professor Stephan Urban of Heidelberg University). The plasmid was transiently transfected into the cells as described previously using PEI. Total RNA was extracted using the FastPure Cell/Tissue Total RNA Isolation Kit (Vazyme, China). One microgram RNA of each sample was reverse transcribed using the HiScript III RT SuperMix (Vazyme, China). The cDNA products were used as templates for PCR to amplify the fragments flanking the A-to-I editing site and the PCR products were subjected to NGS. The primers used in this study were listed in Supplementary Table 4.

To generate HDV dual-fluorescent RNA editing reporter, the original sesRNA sequence in CellREADR vector was replaced by HDV HDAg sequence containing an ADAR-editable translation switch. 5-day post transfection, the fluorescent signal was detected and working efficiency was calculated as described above. The HDV HDAg sequence was listed in Supplementary Table 3.

## Acknowledgements

We acknowledge Dr. Hao Chen (School of Medicine, SUSTech) for his insightful discussions and valuable suggestions that contributed to this study. This work was supported by the National Natural Science Foundation of China (32100766 and 82171416 to R.T., 82270239 to S.S., and 32400478 to T.W.), Guangdong Basic and Applied Basic Research Foundation (2023B1515020075 to R.T.), Shenzhen Fundamental Research Program (JCYJ20220530112602006 and RCYX20221008092845052 to R.T.), Shenzhen Medical Research Fund (A2303039 to R.T. and A2303044 to S.S.), and the China Postdoctoral Science Foundation (2023M731523 to T.W.). We thank Dr. Xibin Lu for his guidance of FACS in SUSTech Core Research Facilities, and Ms. Chunfang Feng for her daily management of the public equipment at School of medicine of SUSTech. We also acknowledge the Center for Computational Science and Engineering at SUSTech for computational resources.

## Author contribution

T.W., J.L., X.L., S.S., and R.T. contributed to the study’s overall conception, design, and interpretation and wrote the manuscript and created the figures with input from the other authors. T.W. designed and conducted the screens of CREDITS and scCREDIT-seq, and validated the screening results, and investigated the molecular mechanisms under the supervision of R.T. J.L. analyzed the scCREDIT-seq and bulk RNA-seq data with guidance from S.S., and R.T. R.L. established the mNeonGreen-tagged *DDX39B* cell line. X.L. designed and conducted the experiments related to RNA editing-based tools. Q.W. and Z.Z. discussed the experiments related to HDV. All authors reviewed and had the opportunity to comment on the paper.

## Supplementary files

Supplementary Figures 1-8 with legends

Supplementary Tables

— Supplementary Table 1: CREDITS screen results analyzed by the MAGeCK-iNC pipeline.

— Supplementary Table 2: sgRNA and siRNA sequences used in this study.

— Supplementary Table 3: Sequences for the *GRIA2*-derived recorder, sesRNAs for CellREADER, arRNAs for LEAPER and HDV HDAg.

— Supplementary Table 4: Primer sequences used in this study.

