## Supplementary Figures for "Multimodal CRISPR screens uncover DDX39B as a global repressor of A-to-I RNA editing"

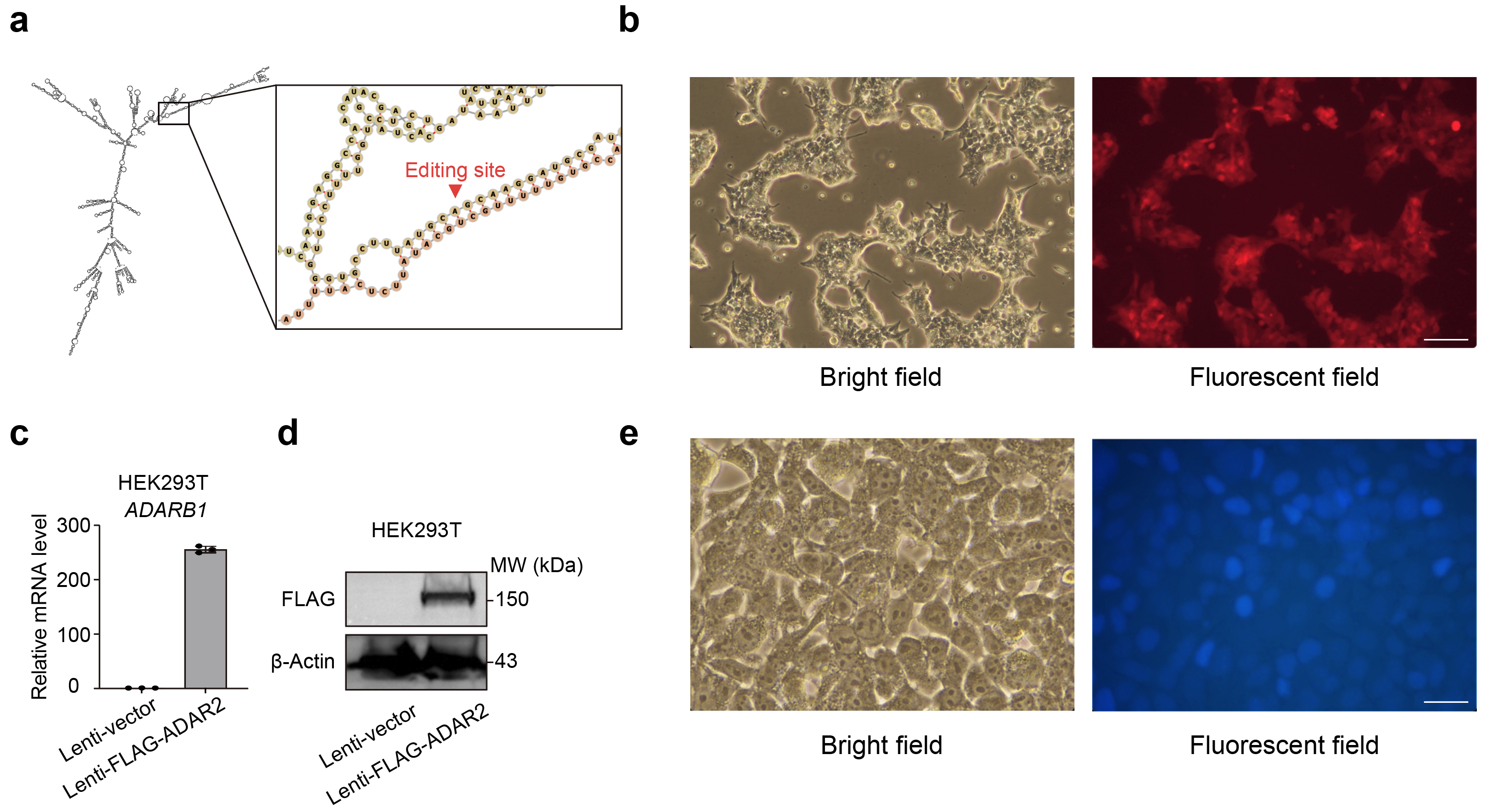

**Supplementary Fig. 1: Generation of the HEK293T cell line stably expressing ADAR2**

**a**, Schematic showing the RNA secondary structure of CREDITS vector and the A-to-I editing site in the recorder (zoomed).

**b**, Representative images of CRISPRi-HEK293T-ADAR2 cells. mCherry signal indicates successful ADAR2 overexpression. Scale bar = 100 μm.

**c**,**d** Measuring ADAR2 (*ADARB1*) expression at mRNA level by qPCR (**c,** mean ± s.d., n = 3 technical replicates) and protein level by western blot (**d**) in HEK293T cells transduced with an empty vector or FLAG-ADAR2 via lentivirus

**e**, Representative images of cells stably expressing the CREDITS vector. Scale bar = 25 μm.

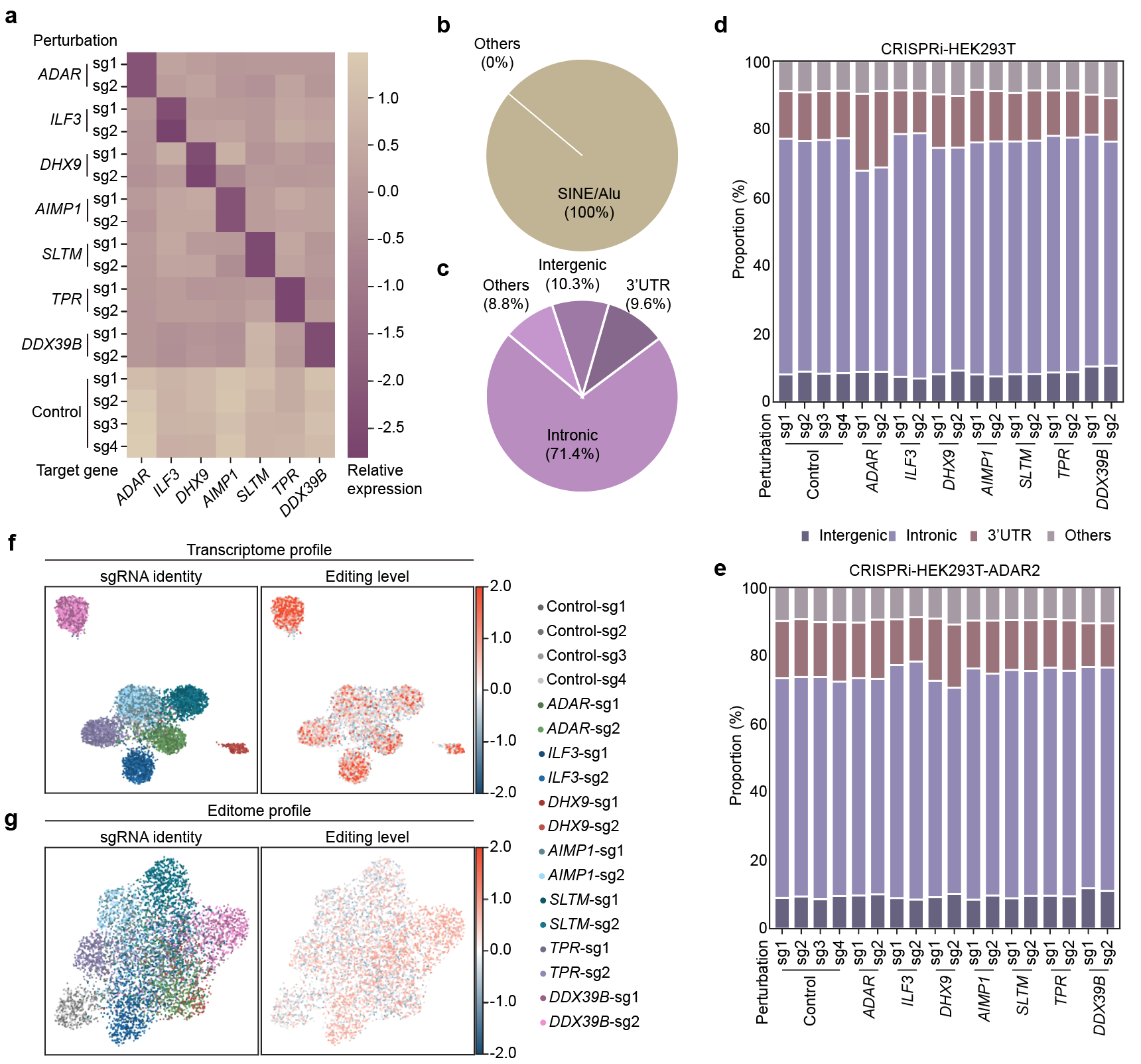

**Supplementary Fig. 2: Characterization of A-to-I RNA editing regulators by scCREDITS-seq**

**a**, Heatmap showing on-target knockdown efficiency for each sgRNA in the scCREDIT-seq screen conducted in CRISPRI-HEK293T-ADAR2 cells.

**b**, **c**, Genomic distribution of high-confident RNA editing sites identified from the scCREDIT-seq screen conducted in CRISPRi-HEK293T-ADAR2 cells.

**d**, **e**, Genomic distributions of RNA editing sites under different perturbations from scCREDITS-seq in CRISPRi-HEK293T (**d**) and CRISPRi-HEK293T-ADAR2 cells (**e**).

**f**, **g**, UMAP visualization of scCREDIT-seq data from CRISPRi-HEK293T-ADAR2 cells following LDA on transcriptome profile (**f**) or editome profile (**g**), color-coded by sgRNAs. The color legend representing each sgRNA was shown (**right**).

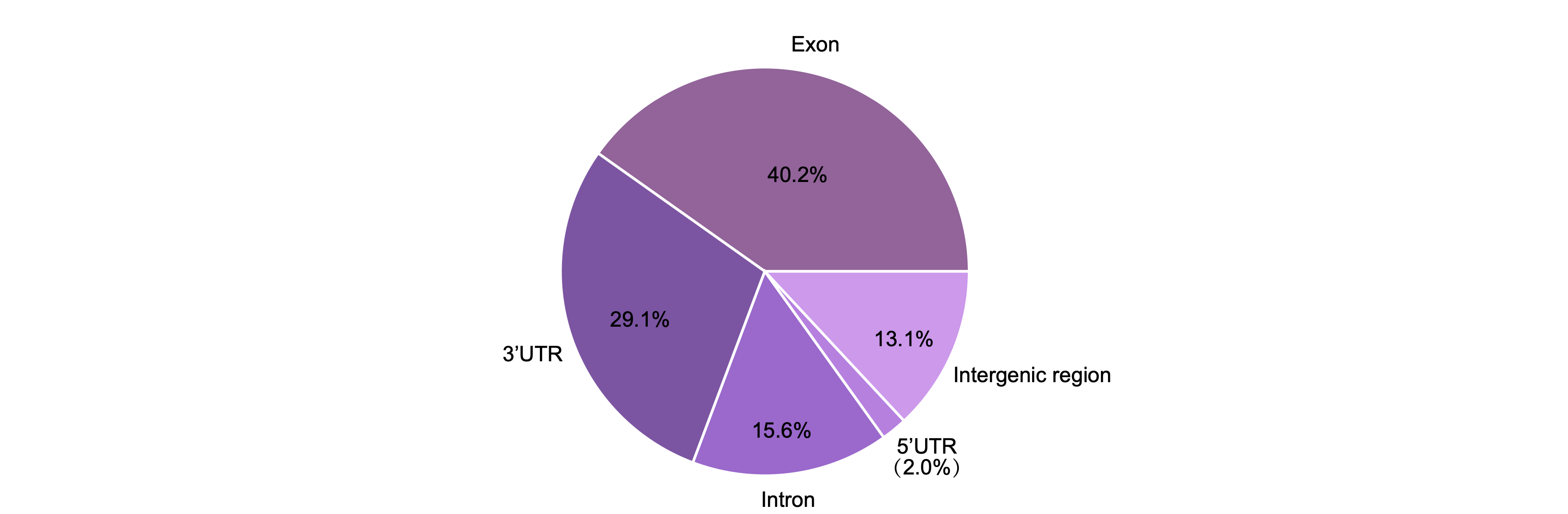

**Supplementary Fig. 3: Reads distribution from scCREDIT-seq**

Genomic distribution of reads from scCREDIT-seq, calculated using RseQC (Version 2.6.5).

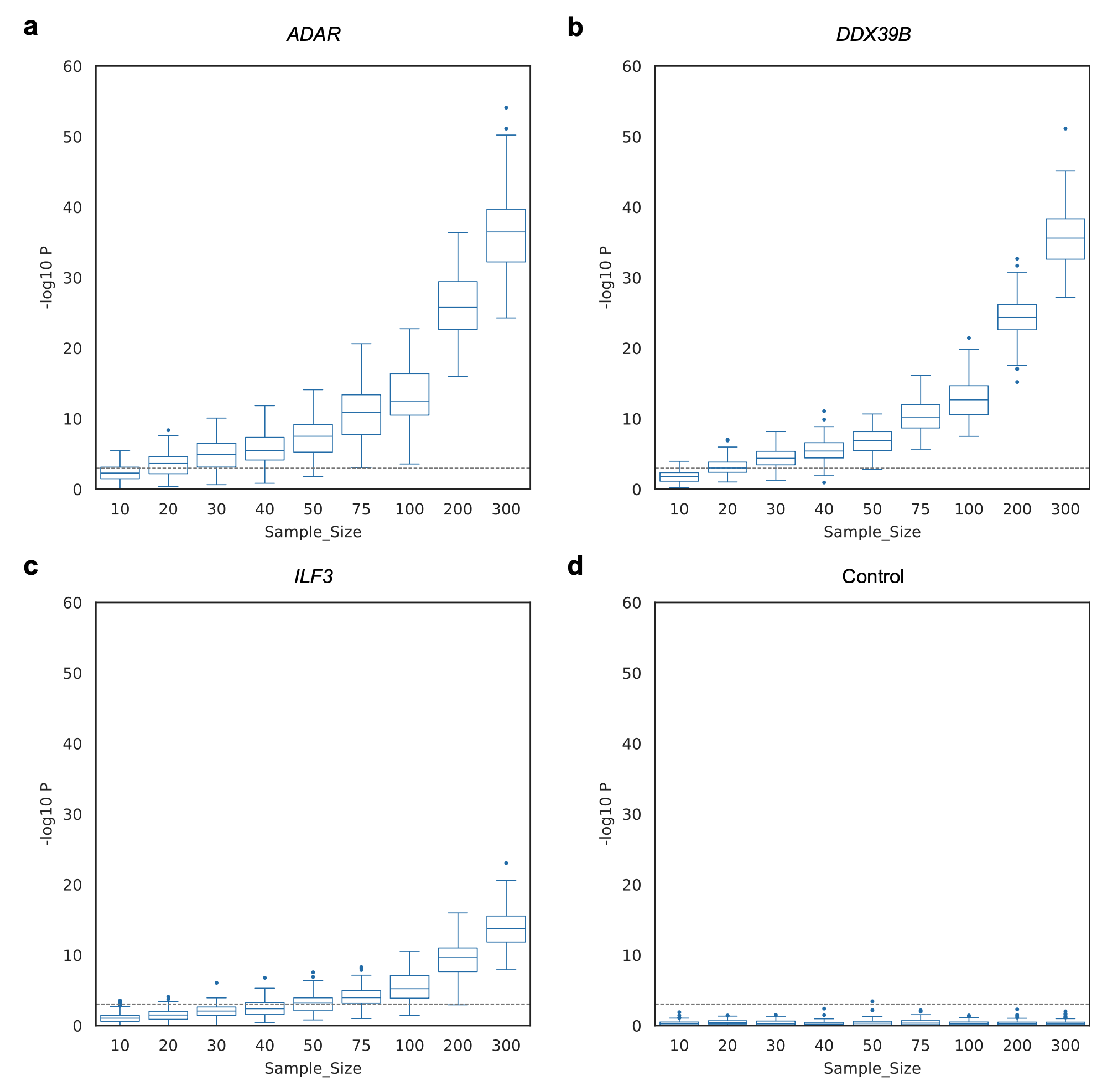

**Supplementary Fig. 4: Bootstrapping analysis of the scCREDIT-seq data**

**a**–**d**, Bootstrapping was performed 100 times with varying sample sizes. Statistical comparisons between specific sgRNA groups and the control group were conducted using t-tests. The distributions of -log10 P values were presented as box plots. The dashed line represents P = 0.05.

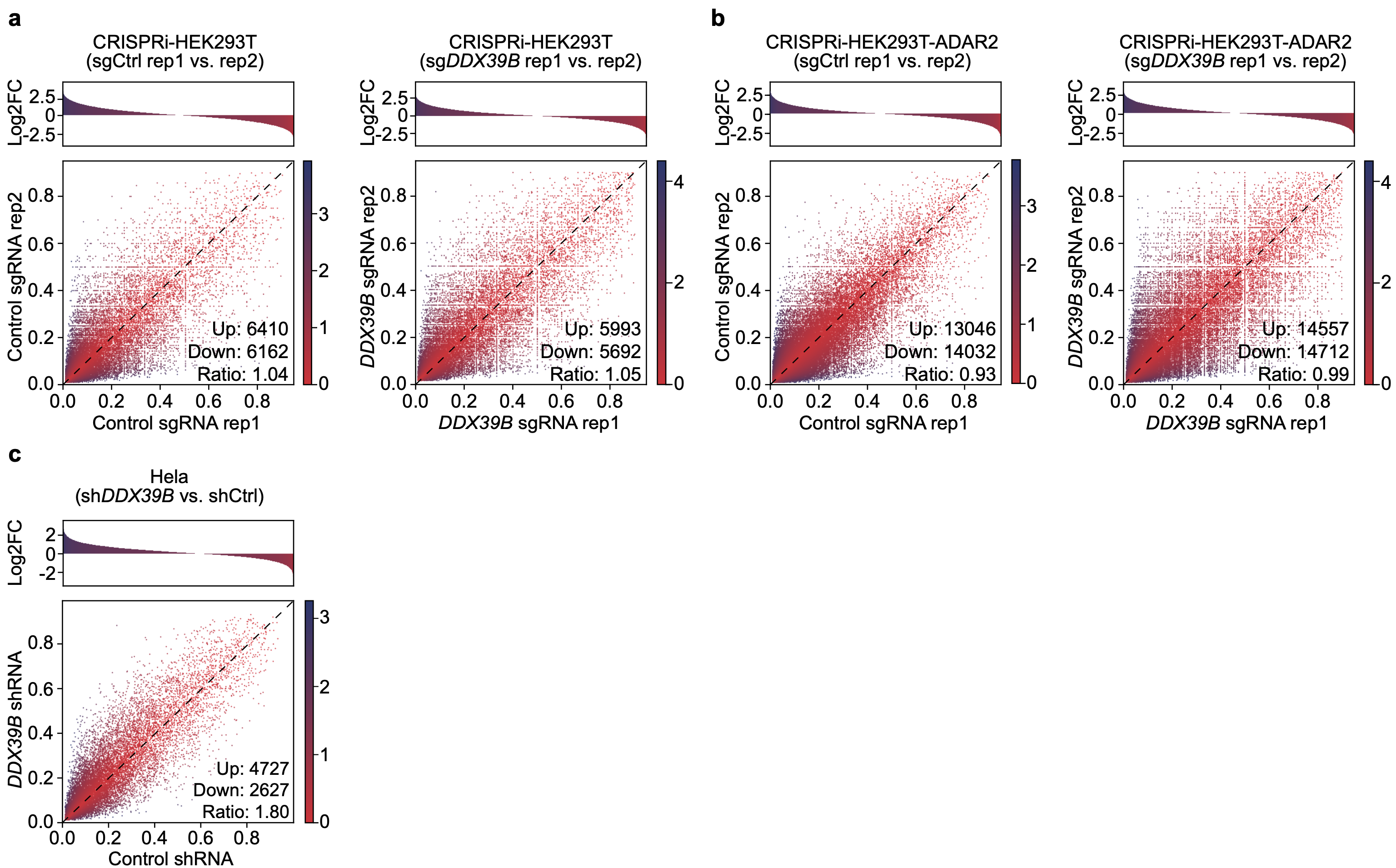

**Supplementary Fig. 5: Bulk RNA-seq analyses of RNA editome**

**a,b,** Differential RNA editing analysis between two biological replicates for control (left) and *DDX39B* knockdown(right) in CRISPRi-HEK293T (**a**) and CRISPRi-HEK293T-ADAR2 (**b**) cells at consensus editing sites. (**Top**) Histogram showing the distribution of log2 fold changes(log2FC) in editing levels (replicate2 / replicate1). (**Bottom**) Scatter plot comparing editing levels between replicate1 (rep1, x-axis) and replicate2 cells (rep2, y-axis), where each dot represents an individual editing site. Colorbar representing log2FC was shown. Numbers of up-edited and down-edited sites (|log2FC| > 0.5) and their ratio were indicated.

**c,** Differential RNA editing analysis for *DDX39B* knockdown in Hela cells. RNA-seq data were obtained from GSE94730^51^.

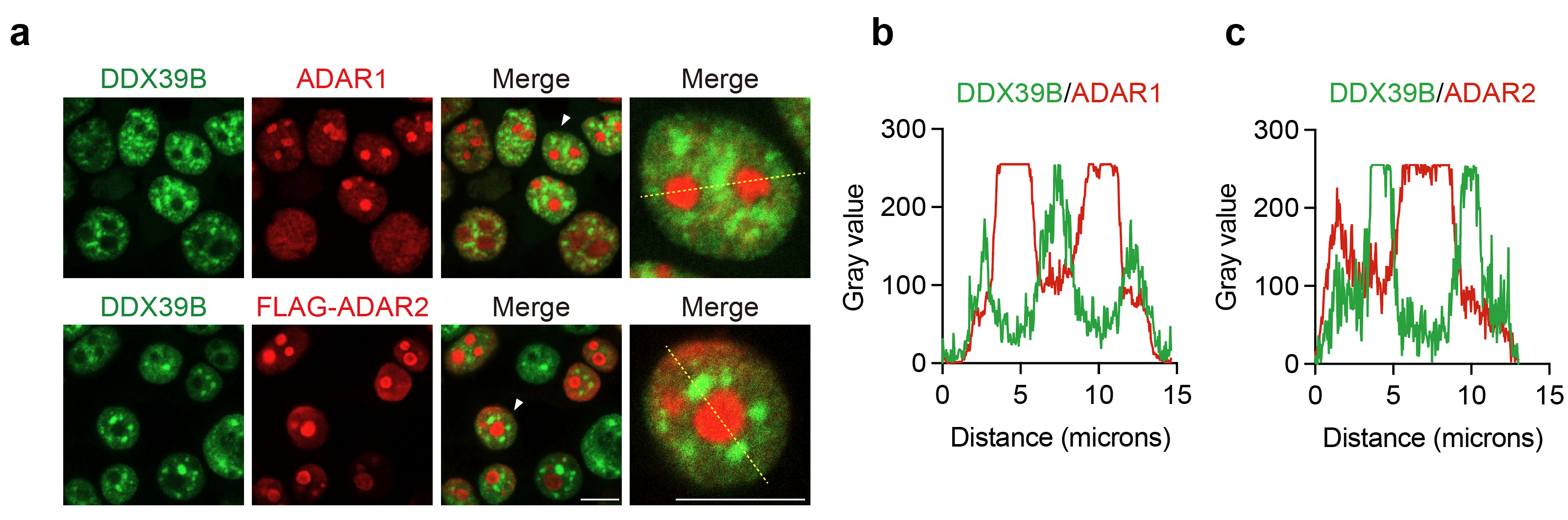

**Supplementary Fig. 6: Localization of DDX39B, ADAR1, and ADAR2**

**a**, Representative immunofluorescence images showing the localization of DDX39B (green), ADAR1 (red), and FLAG-ADAR2 (red) in mNeonGreen-DDX39B HEK293T cells.

**b**, **c**, Quantitative analysis of protein colocalization between DDX39B and ADAR1 (**b**) or ADAR2 (**c**).

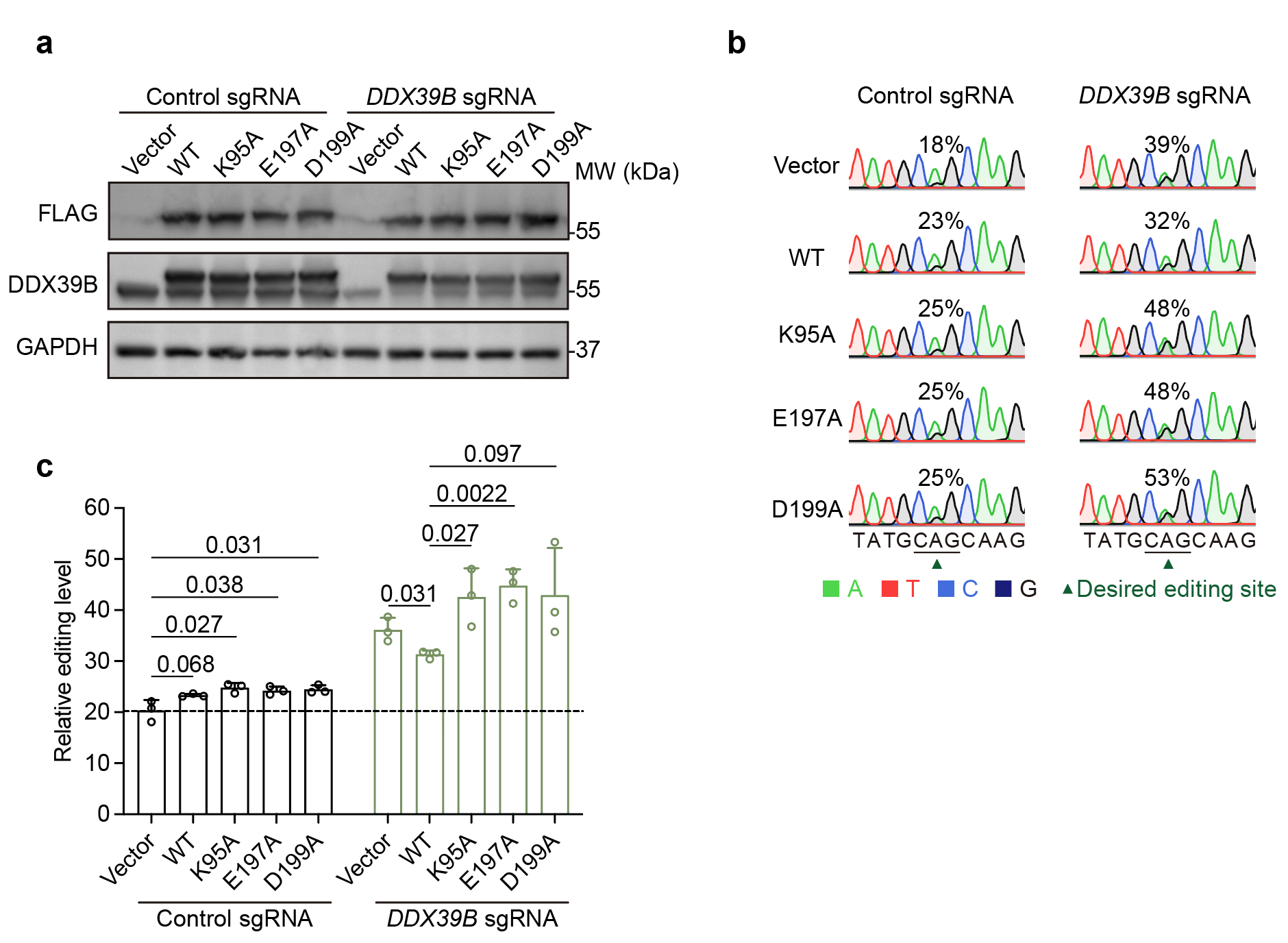

**Supplementary Fig. 7: Rescue experiments**

**a**, Western blot showing the expression of wild-type (WT) and enzymatic-dead mutants of DDX39B in control or *DDX39B* knockdown cells.

**b**, Representative Sanger sequencing electropherograms showing A-to-G conversion in the CREDITS recorder of control and *DDX39B* knockdown cells expressing WT and mutant DDX39B.

**c**, Quantifications of the Sanger sequencing results. P values from unpaired two-sided Student’s t-test were indicated.

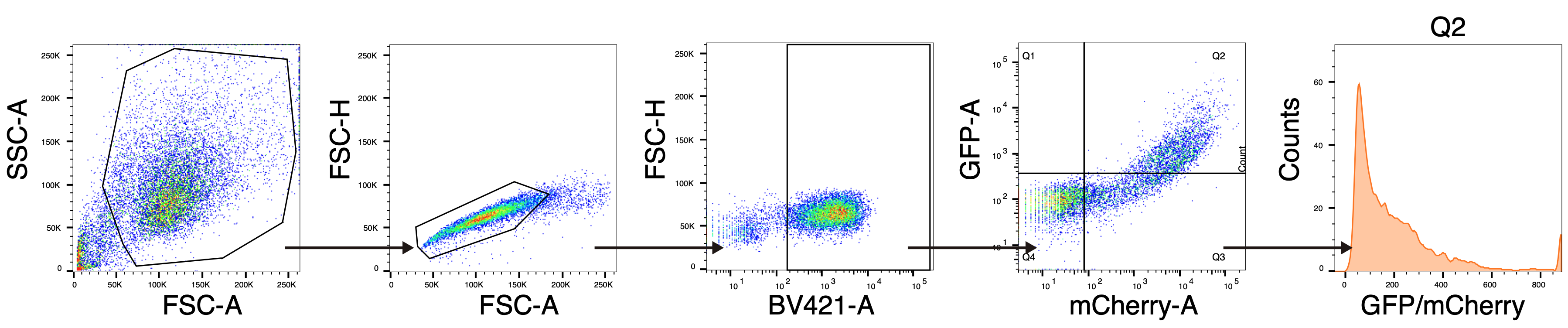

**Supplementary Fig. 8: Gating strategy for flow cytometry analysis**

Representative gating plots for flow cytometry analysis of RNA editing efficiency in CellREADR and HDV genome reporters.
